# Retrospective model utilizing biopsies, granulosa cells, and polar body to predict oocyte competence in bovine

**DOI:** 10.1101/2023.04.23.537973

**Authors:** Dewison Ricardo Ambrizi, Tiago Henrique Camara De Bem, Ricardo Perecin Nociti, Joao Vitor Puttini Paixao, Marcos Roberto Chiaratti, Juliano Ricardo Sangalli, Juliano Coelho Da Silveira, Felipe Perecin, Elisangela Chicaroni de Mattos Oliveira, Jose Bento Sterman Ferraz, Jacinthe Therrien, Lawrence Charles Smith, Flavio Vieira Meirelles

## Abstract

Developmental competence is obtained by a series of morphological and molecular changes during mammalian oocyte growth within the ovulatory follicle. This entails the accumulation of cytoplasmic transcripts that will be used throughout the early stages of development prior to embryonic genome activation, a process known as ooplasm maturation. Furthermore, during follicular growth, epigenetic maturation occurs, which is essential for appropriate embryo development. We believe that transcripts and DNA methylation differ between blastocyst oocytes and those that cleaved but were arrested on day three. We devised a retrospective technique to identify transcripts in oocyte, cumulus, and granulosa cells, as well as DNA methylation connected with oocyte competence, in this work. We dissected and harvested ovarian follicles to achieve this purpose. We extracted and flash frozen the granulosa cells after rupturing them. The oocytes were put in maturation media droplets, and the cumulus cells and polar body were removed and kept the following day. To prevent spermatozoon interference, we chemically activated the oocytes and tracked their development (until they reached the blastocyst stage). We went back to their biopsies, cumulus cells, and polar bodies and did RNA-seq (biopsies and cumulus) and single polar body WGBS (polar bodies) when we collected the results 7 days later, i.e. 1-) embryos that cleaved but stopped development (termed CL) or 2-) embryos that cleaved and progressed until the blastocyst stage (termed BL). Additionally, after transcriptome results from oocyte-biopsy and cumulus cells, the granulosa cells from their individual oocytes were sequenced as a noninvasive strategy. This study is a follow-up to our previous work, “**Assessment of Total Oocyte Transcripts Representation in bovine Using Single Ooplasm Biopsy with High Reliability.**” Following sequencing, we discovered that the two groups, BL x CL, were transcriptionally different in granulosa and biopsy samples, although cumulus cells were a poor predictor of oocyte competence. By analyzing the differentially expressed genes, we discovered multiple genes and pathways related to oocyte competency, demonstrating the efficacy of our method. Despite no change in morphology, these alterations in pathways and genes show that the oocyte CL group was transcriptionally and epigenetically delayed, with ferroptosis and necroptosis processes activated. The oocyte BL group demonstrated numerous molecular signaling, oocyte meiosis, GnRH signaling, G-protein cascade, and RNA stability pathways. In network analysis, we discovered *GNAS*, an imprinted gene and one of the most important essential genes. The transcripts from granulosa cells confirm the oocyte results. Nonetheless, transcriptional variations in granulosa cells were far greater than those in oocytes (97% × 34% variance), implying a completely distinct transcriptome in the follicular niche. In terms of the WGBS, we discovered differentially methylated areas linked with oocyte competency, as well as transcriptome results confirming the structure’s ability to predict outcome. These findings might be beneficial in clinical settings for those undergoing infertility therapy. In the oocyte, we discovered a complex transcriptional and epigenetic regulation network. Furthermore, mature cumulus transcription produced information that differed from the true content of the MII oocyte and granulosa cells before maturity. Our findings underscore the significance of maternal transcripts and epigenetic maturation in early parthenogenesis, as well as the use of granulosa cells as early indicators of competence.

## INTRODUCTION

The mammalian ovarian reserve is a large but finite reservoir of primordial follicles holding oocytes in diplotene that govern female reproductive potential (McGee & Hsueh, 2000). Despite the fact that the ovarian reserve contains millions of oocytes, only a small percentage of them will be chosen to mature during folliculogenesis and gain the potential to produce a viagle gamete over the curse of a female’s reproductive lifespan (Duan et al., 2019). During the acquisition of developmental competence, oocyte growth is closely associated with follicular growth and involves progressive structural and molecular changes that include, at the follicular level, the accumulation of follicular fluid and an increase in the number of somatic cells and, at the oocyte level, the accumulation of transcripts to be used during early development and acquisition of correct epigenome (Labrecque et al., 2015; Wei et al., 2019; Trebichalská et al., 2020).

According to Matoba et al. (2014), there exists a correlation between the composition of follicular fluid and the development of embryos, although no such correlation has been observed with respect to hormone levels. The acquisition of accurate oocyte imprinting has been found to be significantly associated with embryo development, as demonstrated in mice with regards to genes such as H19, Gtl2, and Peg10 (De Santis et al., 2005; Fan et al., 2002; Wei et al., 2019;). The process of follicle development and oocyte maturation is associated with the transcriptional regulation of mural-granulosa cells (Alam & Miyano, 2019). Extensive research has been conducted on a dynamic interaction between somatic cells and oocytes in mammals, such as humans and cows, as reported by Liu et al. (2018). Various stimuli can elicit a response from granulosa and granulosa-cumulus cells, leading to alterations in the microenvironment of the follicle. The accumulation of transcripts, lipids, and other molecules in the oocyte is facilitated by various mechanisms such as gap junctions, transzonal projections, and extracellular vesicles, which are mediated by granulosa-cumulus cells (Sutton et al., 2003; Macaulay et al., 2014; del Collado et al. 2017). Significantly, research has established a correlation between granulosa cell transcription and oocyte competence, and has suggested that granulosa cell transcription could serve as a promising biomarker in various species including mice, cows, pigs, and humans (Li et al., 2008; Uyar et al., 2013; Mazzoni et al., 2017; Meng et al., 2020). These studies provided a comprehensive assessment of oocyte contents using granulosa-cumulus transcripts. Whilst these studies have established an association between granulosa-cumulus cells and oocyte competence, they have not established a direct correlation between specific ooplasm transcripts and oocyte quality.

The capacity to restart meiosis and complete the first meiotic division with the ejection of the first polar body and to arrest at the metaphase stage of the second meiotic division, also known as MII, constitutes the acquisition of oocyte competence (Conti & Franciosi, 2018). Apart from meiotic events, cytoplasmic components related to oocyte competence have already been discovered (Mamo et al., 2011; Watson, 2007). The oocyte accumulates important molecules in the cytoplasm during follicular development until the conclusion of the germinal vesicle phase, which can be connected to competence (Brevini Gandolfi & Gandolfi, 2001). The accumulation of such factors, which includes RNAs, is associated with the ability to resume meiosis until MII, allows conditions for normal fertilisation by avoiding polyspermy, and enables the fertilised zygote to cleave and coordinate initial development until the major embryo genome activation (EGA), which occurs between the 8 to 16 cells in bovine (Albertini et al., 2003; Brevini Gandolfi & Gandolfi, 2001; Conti & Franciosi, 2018; Yang et al., 2020). As a result, maternal RNAs that accumulate in the oocyte are required for chromatin reorganization and transcription factor translation (Eckersley-Maslin et al., 2018; Østrup et al., 2012). Despite attempts to estimate oocyte potential using mural granulosa, cumulus cells, extracellular vesicles from follicular fluid, and other follicular niche components, results are conflicting and unable to correlate precisely which molecules in the ooplasm are responsible for the oocyte’s ability to develop to the blastocyst stage (Gad et al., 2022; Martínez-Moro et al., 2022).

Beyond the RNA world, epigenetic maturation is an important event that must occur in the oocyte, and it is directly related with the coordination of global events in the oocyte genome (Yuan et al., 2020). Correct methylation of *Snrpn, Peg3, and H19*, for example, is required for blastocyst development in mice but is imbalanced in bi-maternal embryos (Wei et al., 2019). The use of the first polar body to examine the oocyte methylation genome using WGBS is practical and has a strong correlation (0.92), allowing it to be utilized to research methylation without destroying the oocyte (Yuan et al., 2020).

Our hypothesis posits that the follicular niche is associated with a molecular signature that indicates the quality of the oocyte. The present investigation employed a retrospective methodology to identify the MII transcripts in the ooplasm and the adjacent granulosa cells (cumulus), in addition to assessing the methylation levels in the first polar body. In addition, we also investigated transcripts obtained from granulosa cells that were collected during the GV phase. The experimental design is presented in Figure 1A centered on the capacity of the oocyte to attain the blastocyst stage in vitro subsequent to parthenogenetic activation. The utilization of this invasive technique permits the evaluation of MII RNAs in oocytes while minimizing interference with their developmental potential and ensuring a comprehensive representation of the entire oocyte (as per the methodology publication). Following normalization, transcripts were found to be associated with oocyte and follicular competence. The identification of key pathways involved in the acquisition of developmental competence was facilitated through the utilization of differentially expressed genes (DEG), hub genes, and exclusive genes. In addition, the examination of transcripts in granulosa cells obtained through follicular aspiration provided an evaluation of the follicular environment on the day of collection, as well as a comparison of the biological processes operating concurrently with developmental competency.

**Figure 1:**
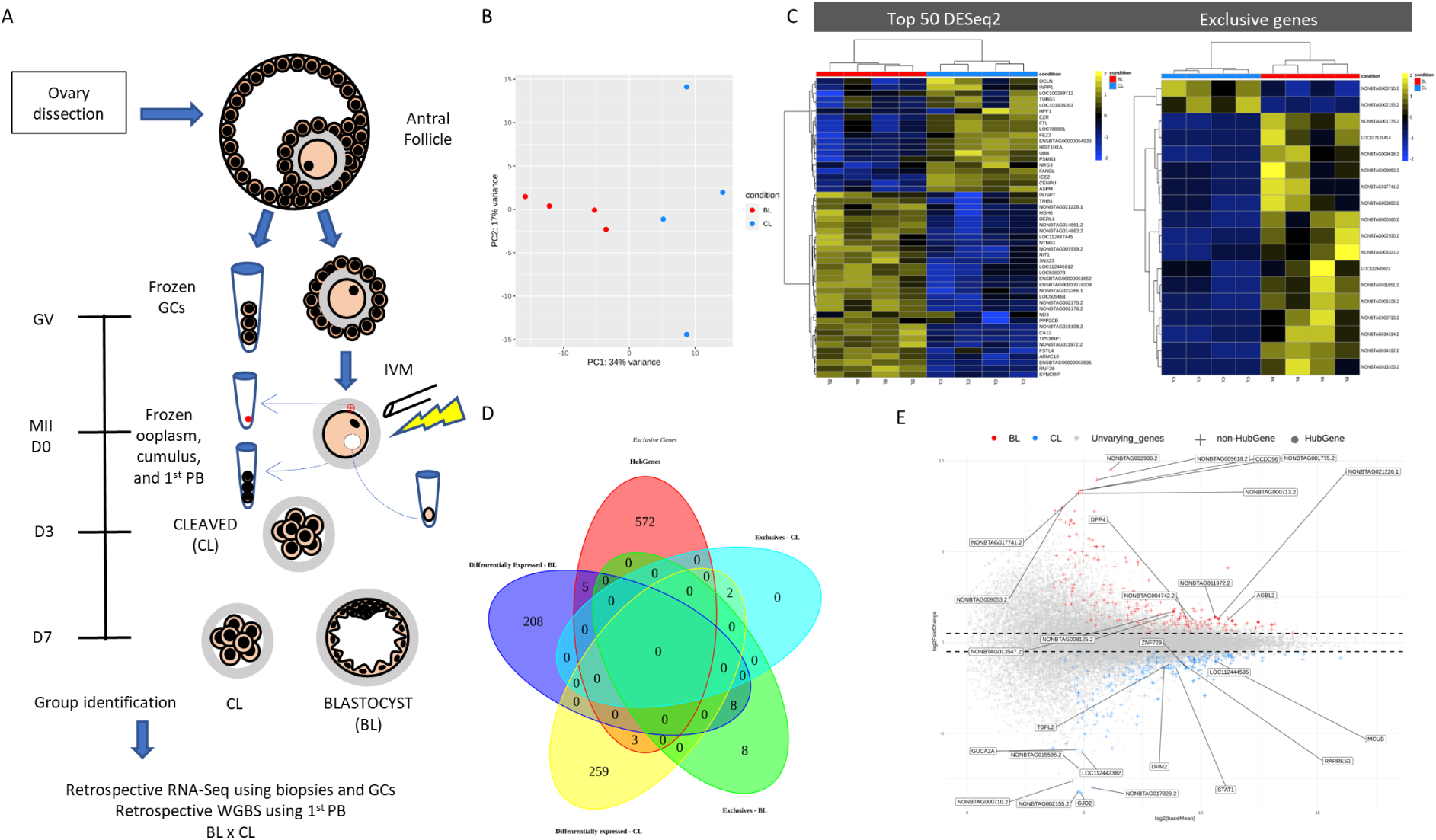
Transcriptome analysis of oocytes. A - Retrospective model design for group identification (BL x CL) using individual oocyte maturation and cultivation. The cleavage (CL) rate was determined on day three, whereas the blastocyst (BL) rate was determined on day seven. For biopsies, cumulus, and granulosa cells, BL and CL were then sequenced using the Single-cell RNA-Seq method. Single WGBS was used for PB. B - PCA demonstrating 34% variation in PC1 and 17% in PC2. C - Heatmap of the 50 DEGs with the greatest difference between groups; exclusive genes were those that were present or absent in one of the groups (right heatmap). D - Venn diagram illustrating a logical relationship between sets of significant gene pick-up methods. Smearplot to show the main differences between the BL and CL groups. The BL group was symbolized by red, whereas the CL group was represented by blue. We used spheres to describe hubgenes and addition symbols to identify non-Hubgenes. The first seven Hubgenes and DEGs in each group were labeled based on log2FoldChange value.

## RESULTS

Samples used in this study were obtained exclusively of dissected follicles raging from 4 to 5-mm containing COCs (cumulus-oocyte complex) and granulosa cells with superior morphology. After in vitro maturation, COCs were denuded of cumulus cells and checked for the presence of the first polar body (1^st^ PB). Only MII oocytes with normal morphology were biopsied, and then activated. Oocytes were checked for cleavage on day three and blastocyst development on day seven. The development rates of each of the routines can be visualised in Table 1. Using a single-cell RNA-seq approach (SMART-Seq2), we generated an average of 10 million paired-end reads per biopsy sample. Granulosa samples of the respective biopsies were sequenced using the RNA-Seq approach.

**Table 1.**
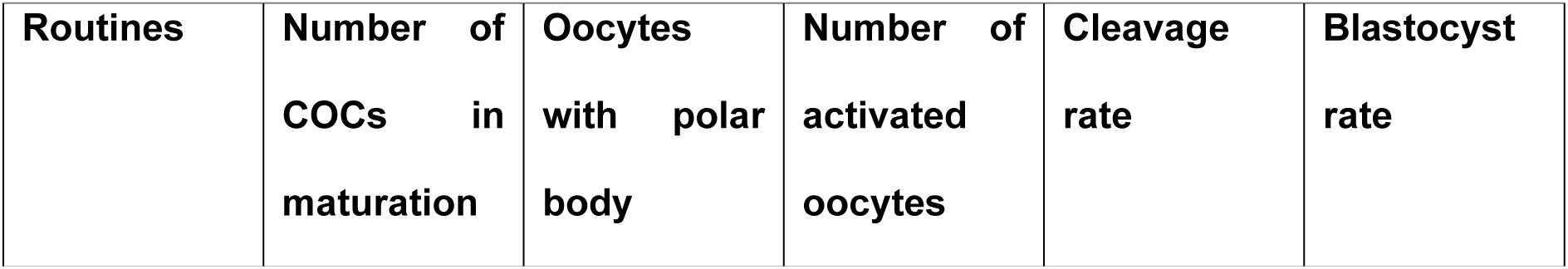
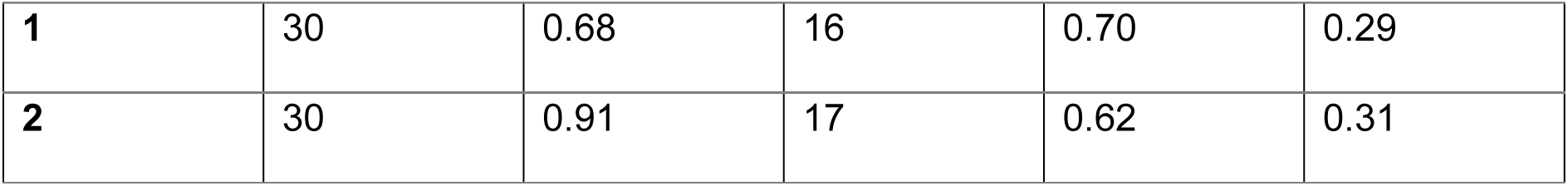
Development rates of the oocytes cultured individually in this study.

| Routines | Number of COCs in maturation | Oocytes with polar body | Number of activated oocytes | Cleavage rate | Blastocyst rate |
| --- | --- | --- | --- | --- | --- |
| 1 | 30 | 0.68 | 16 | 0.70 | 0.29 |
| 2 | 30 | 0.91 | 17 | 0.62 | 0.31 |

### Ooplasm critical pathways associated with developmental competence

To validate the use of the retrospective approach, a preliminary study showed a ∼99% similarity between the transcriptomes of the oocyte biopsies and the remaining oocyte. Biopsies from oocytes that cleaved and then either arrested or developed to the blastocyst stage after parthenogenetic activation produced an average of ten million reads per sample. After DESeq2 normalization, we identified 27555 genes and, using PCA analysis to assess total variance, we showed a 34% variance in PC1 and a clear clustering of the samples according to competence (Figure 1B). All comparisons were performed using BL against the CL group being designated as upregulated when the gene was a DEGs (differentially expressed gene, absolute log2FoldChange > 0.5 and adjusted p < 0.1) in BL group, while downregulated was used to CL DEGs. A heatmap using all normalised genes also showed the complete clusterisation (ward D2) of the groups according to competence (Figure S1A). We further examined in a reduced heatmap the fifty genes with the highest variation between groups (Figure 1C), which enabled the identification of some common processes. Genes with no symbol nomenclature were blasted using ensemb (*Bos taurus, Homo sapiens*), and NCBI (*Bos indicus, Homo sapiens*) browser blast tool in order to identify the most similar homologous transcripts. In an overview, we could concatenate these genes to some process, such as blastocyst hatching process (*NONBTAG014862.2* | *NONBTAG014861.2* = *SMIM14, FST4*), biomarkers associated with oocyte quality (lncRNA - *ENSBTAG00000053835*), RNA stability (*SYNCRIP, NONBTAG011972.2*), regulation of cell growth (*PPP2CB, NONBTAG002176.2* | *NONBTAG002175.2* = *MOB*, *LOC508073* | *LOC112445812* = melanoma-associated antigen 10-like, *LOC112447445* = *NDFIP1*, *NONBTAG007959.2* = *S100P*, *RIT1, TP53INP1*), negative regulation of transcription (*ENSBTAG00000019009* | *ENSBTAG00000051652* = *MAGEA13P*, *NONBTAG021226.1* = *SETD5*), signal transduction (*RNF38*), metabolism regulation (*ARMC10*, *CA12, ND3, LOC505468 = CYP2C88*), cytoskeleton organization (*NONBTAG015108.2* = *SEPTIN5*, *NONBTAG022266.1 = TMED7, NTNG1*), degradation of the unfolded protein (*DERL1*), and DNA mismatch repair (*MSH6*), all linked to BL group. On the other hand, in the CL group, we can see a conversion for chromatin organisation (*HPF1, LOC100299712 = ZNF665, HIST1H1A, ENSBTAG00000054933 = H4C2, CENPU, ICE2, TUBG1*) neuronal development (*NGR3, FEZ2, ASPM TUBG1, LOC101906393*), ubiquitination process (*PSMB3, UBB, FANCL*), iron homeostasis (*FTL, LOC788801*), communication (*EZR, OCLN*), and phosphatidylinositol (*INPP1*).

**Figure S1.**
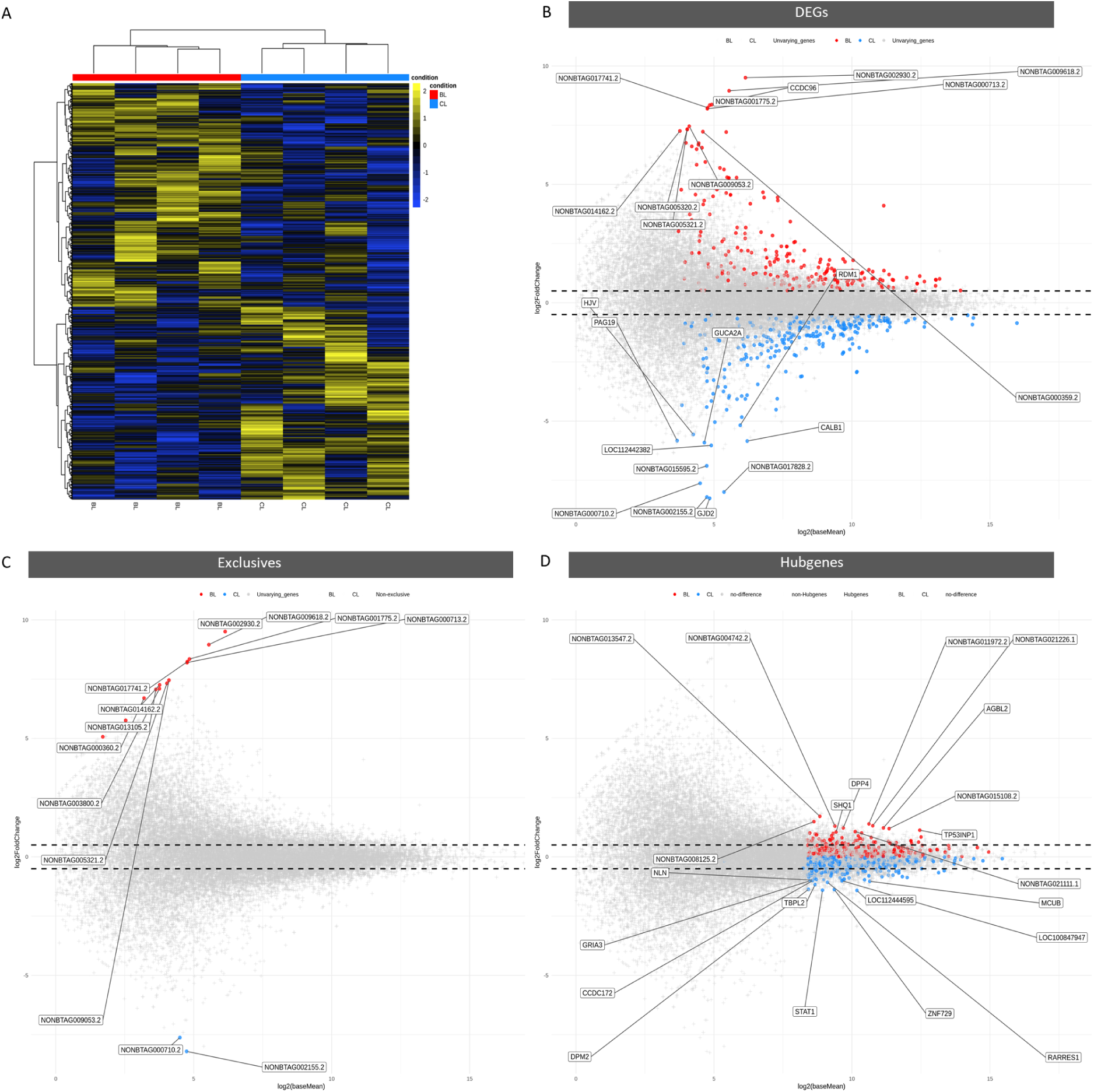
Clusterization and different dispersion of DEGs, exclusives and Hubgenes. (A) Heatmap using total DESeq2 normalized genes (27555) showing a clear clusterization between groups BL and CL. Smearplot for DEGs (B), exclusive (C), and Hubgenes (D).

Genes found in one group and not in the other were also investigated to improve the biological processes (TPM >= 1 in 70% of samples in one group and precisely equal to 0 in the other) enabling the identification of eighteen exclusive genes, i.e., 16 for BL and 2 for CL group (Figure 1C). By contrasting the BL and CL groups we identified 486 DEGs (221 - BL, 264 - CL). Moreover, using the CeTF R package, we identified 580 Hubgenes (LFC > 2 and p-adj < 0.05) among which eight were both Hubgene and DEGs simultaneously (BL = *NONBTAG015108.2, NONBTAG021226.1, NONBTAG011972.2, NONBTAG013547.2, TP53INP1; CL = LOC101907041, MCUB (ENSBTAG00000012995), ZNF729*), and ten simultaneously as DEGs and exclusives (BL = *NONBTAG002930.2, NONBTAG009618.2, NONBTAG001775.2, NONBTAG017741.2, NONBTAG000713.2, NONBTAG009053.2, NONBTAG005321.2, NONBTAG014162.2;* CL = *NONBTAG002155.2, NONBTAG000710.2*). These classificatory differences are presented in a Venn diagram (Figure 1D) and a the dispersion (log2 baseMean x log2 FoldChange) smearplot (Figure 1E). We labelled the seven genes with the highest log2FoldChange in both groups for DEGs and Hubgenes. For the BL group: *NONBTAG002930.2, NONBTAG009618.2, CCDC96, NONBTAG001775.2, NONBTAG017741.2, NONBTAG000713.2, NONBTAG009053.2, NONBTAG013547.2, NONBTAG008125.2, NONBTAG011972.2, NONBTAG021226.1, NONBTAG004742.2, AGBL2, DPP4*; for the CL group: *GJD2, NONBTAG002155.2, NONBTAG017828.2, NONBTAG000710.2, NONBTAG015595.2, LOC112442382, GUCA2A, LOC112444595, STAT1, ZNF729, DPM2, TBPL2, RARRES1, MCUB (ENSBTAG00000012995)*. A smearplot for DEGs, exclusives, and Hubgenes can be seen in Supplementary Figure 1B-D.

After identifying the DEGs, exclusives, and Hubgenes, to explore in a general view, we constructed a concatenated gene list to investigate critical pathways (Supplemental file 1) and performed cluster analysis for Kegg (Figure 2A). In the BL cluster, 122 genes composed pathways such as MAPK, Relaxin, Kaposi sarcoma-associated genes, Autophagy, Oxytocin, Chemical carcinogenesis - reactive oxygen species, Toll-like receptor, GRH, FoxO, lysosome and pathways related to the nervous system were among the pathways in this cluster. When considering only DEGs, butanoate metabolism, cGMP-PKG-signalling, Oocyte meiosis, and TGF-beta were also identified (Figure S2).

**Figure 2.**
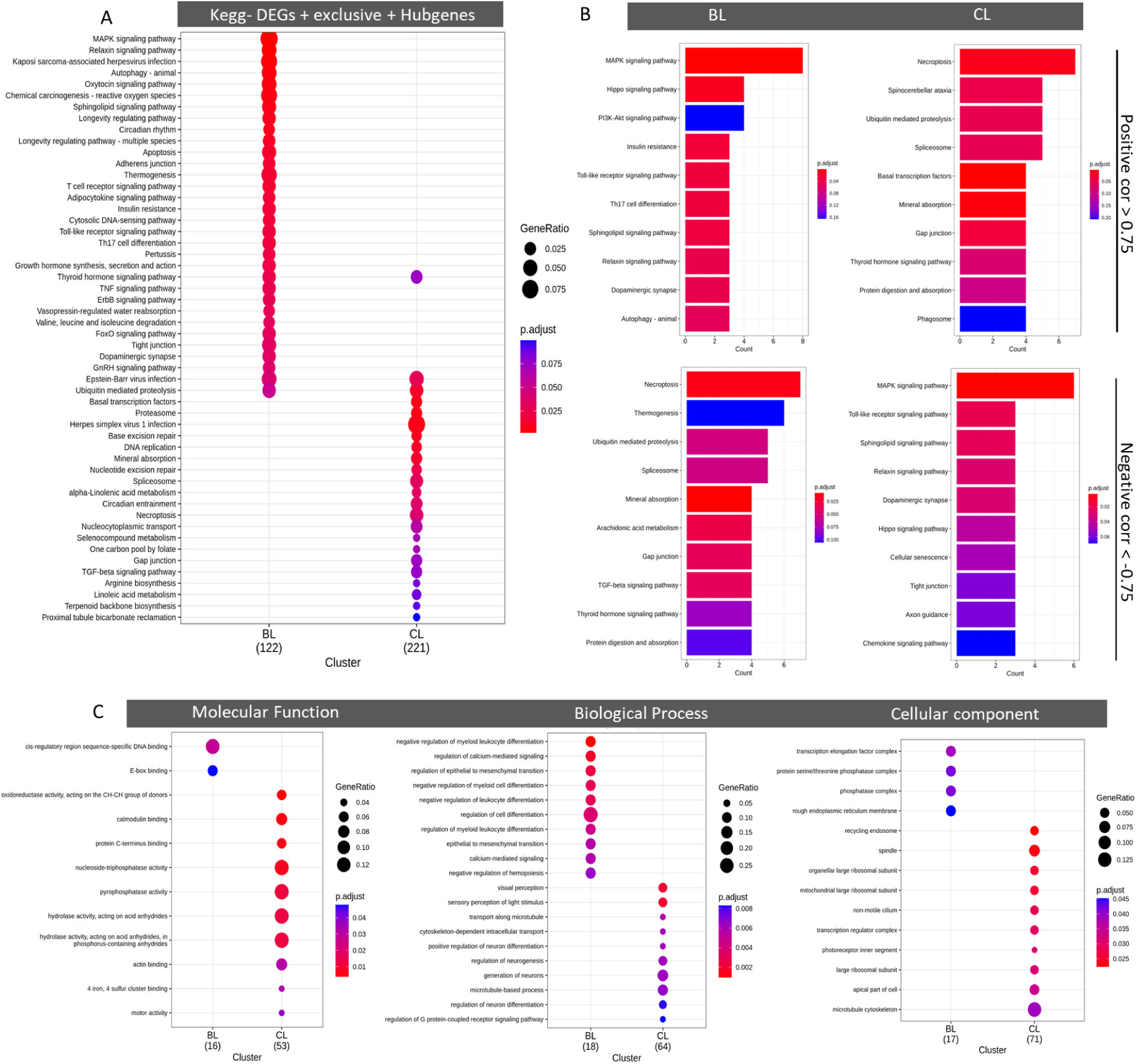
Critical pathways differences linked to oocyte competence. A - A total of 122 and 221 were found in compare cluster analysis respectively for the BL and CL groups. B - Critical pathways highly correlated with genes found in overlapp as DEGs, exclusives and Hubgenes (sixteen genes for BL and five for CL) were used. We filtered absolute Spearman correlation values > 0.75 and with p-value < 0.05. C - Gene ontology pathways constructed using DEGs + exclusives + Hubgenes. They demonstrate the activity that gene product performs (Molecular function), the specific objective that those genes can be programmed to achive in the organism (Biological process), and where the product can have a function (cellular component)

We constructed the Gene ontology (GO) analysis for molecular function (MF), biological process (BP), and cellular component (CC) using the merged gene list. This analysis identified transcription regulatory pathways (MF - cis-regulatory sequence-specific DNA binding, E-box binding) with strict regulation of cell differentiation and calcium signalling (BP), with possible action in transcription elongation factor, phosphatase complex and rough endoplasmic reticulum membrane for the BL group. The CL GO analysis identified hydrolase/phosphatase activity, actin binding (MF), and neuronal development (BP); for the CC, spindle, apical part of the cell, and microtubule cytoskeleton can indicate where the product of those genes can act (Figure 2C).

**Figure S2.**
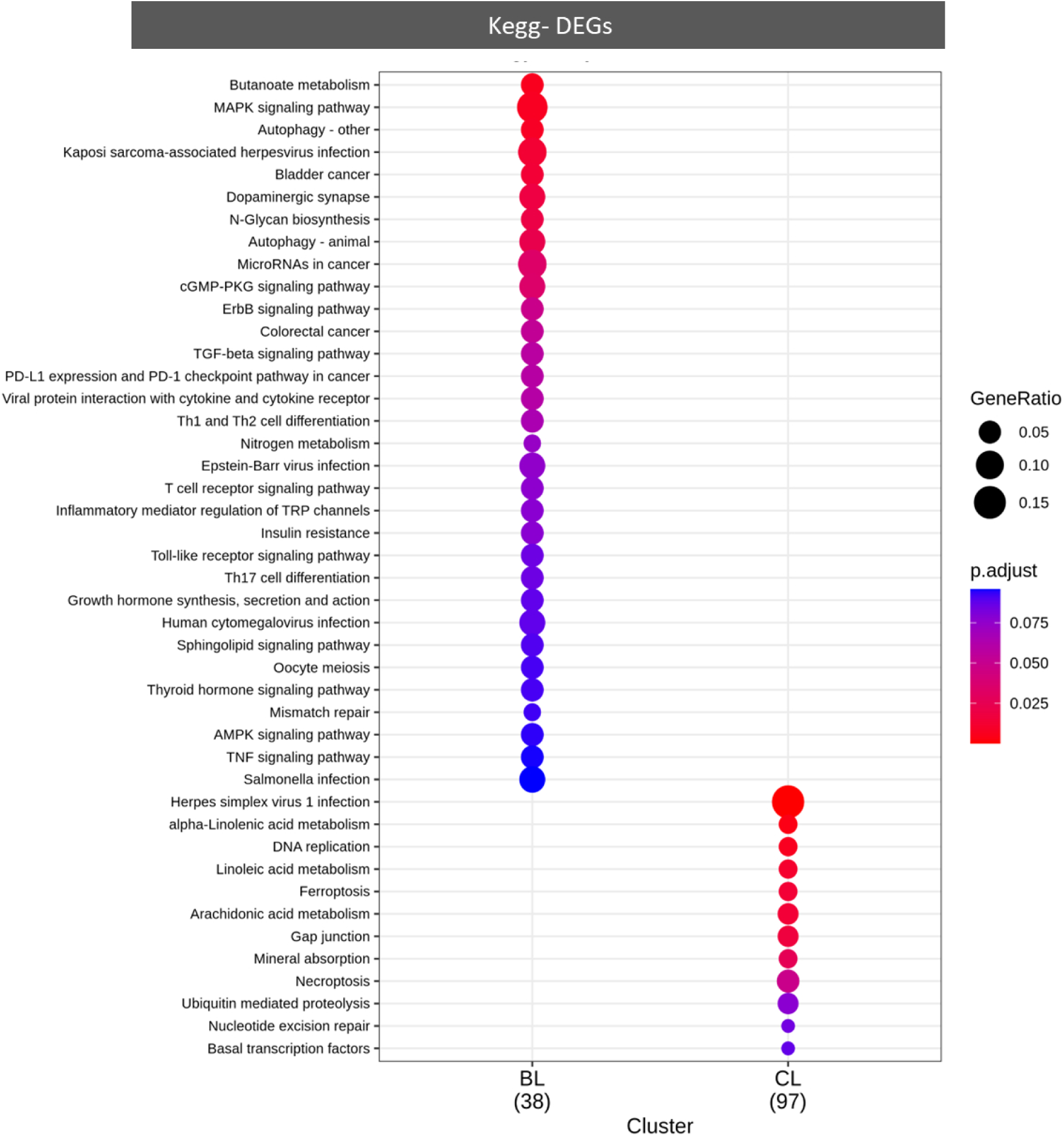
Main pathways using DEGs. (A) and network Kegg analysis using merged gene list (B). The network analysis identified the most important genes and pathways in the graphic center.

MAPK and Sphingolipid signalling pathways were more activated in the BL group and presented genes associated with anti-apoptosis (NFkB, c-Myc, Ras, ERK1/2, MEK1/2, and PKCL), and G-protein subunit S (Gs/olf). On the other hand, the 221 genes in the CL cluster participated in the Thyroid hormone, Epstein-Barr virus infection, Ubiquitin mediated proteolysis, Basal transcription factors, Proteasome, DNA-replication, Nucleotide excision repair, TGF-Beta. Herpes virus infection, Base excision repair, alpha-Linolenic metabolism, and Necroptosis pathways; ferroptosis was found using only DEGs. Herpes simplex virus 1 infection and Necroptosis were found in the CL group with genes associated with apoptosis (BAX, BCL-2 and p53, FTH1, cPLA2), spliceosome, and ubiquitination (UCHL1, UBC6/7), and axonal transport defects (TUBB) (Figure S3).

**Figure S3.**
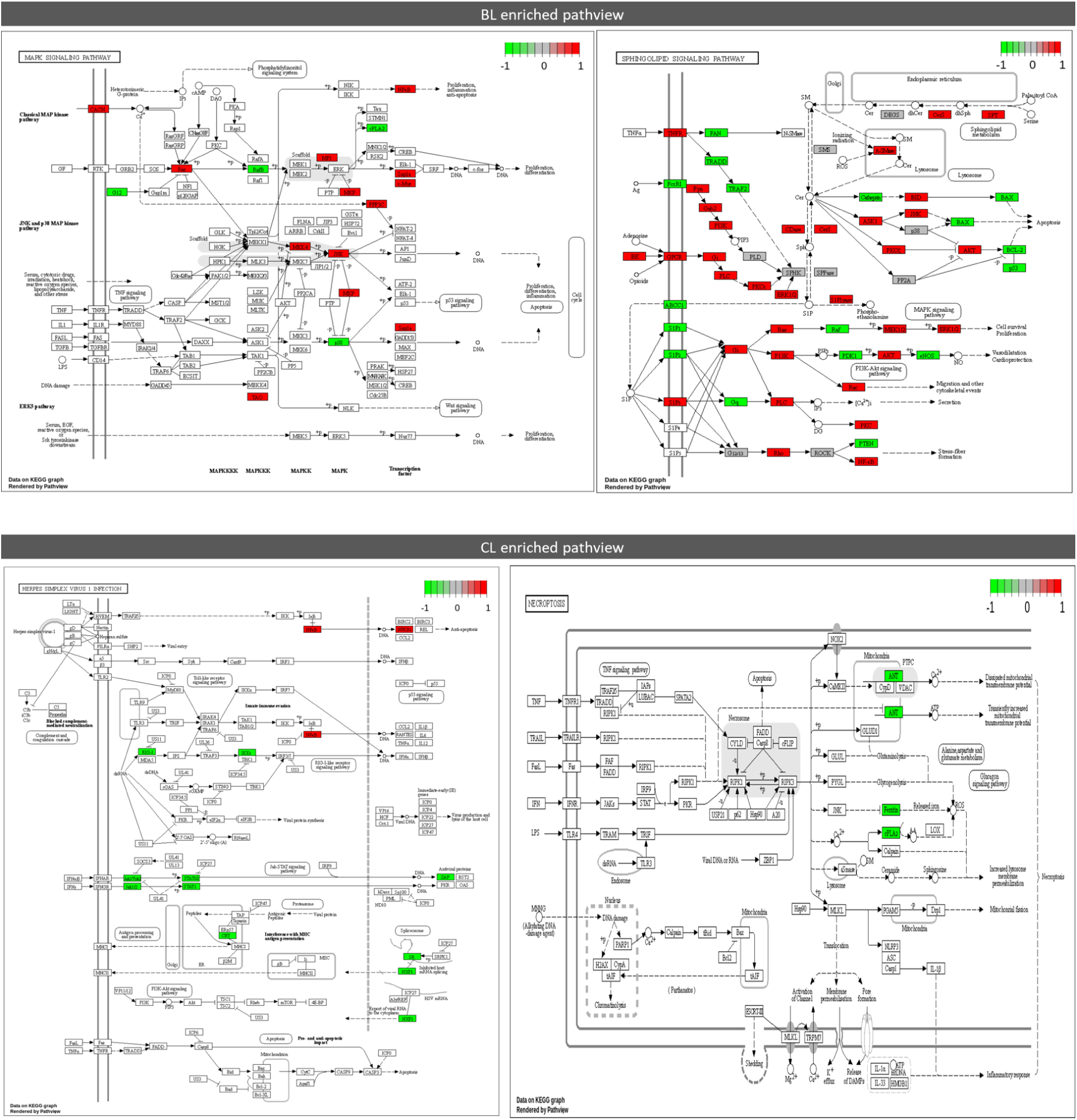
Pathview visualization for main pathways enriched for BL and CL groups. Red color represents the BL group while green color represents the CL group.

Next, as aforementioned about genes categories, we investigated the co-expression correlation network of genes in the overlap of DEGs and Hubgenes or DEGs and exclusives. We applied Spearman correlation using all DESeq2 normalised genes to explore which genes are positively or negatively co-expressed with overlapped genes for each group. The genes *NONBTAG011972.2, NONBTAG013547.2, NONBTAG015108.2,* and *NONBTAG021226.1* in the BL group and ZNF729 in the CL group were blasted (*Homo sapiens, Bos taurus, Bos indicus*), resulting in respectively of *TENT5C* (mRNA stability), *KIAA1147* (exchange of GDP to GTP), *SEPTIN5* (cytoskeletal organisation), and *ZNF729* (predicted to enable DNA transcription factor activity), suggesting the biological functions for these genes. For both groups, we filtered only absolute correlations higher than 0.75 with a p-value smaller than 0.05 resulting in 4258 genes for BL and 1601 for CL (Supplemental file 2). The BL positive correlated genes enriched for practically same pathways found in CL negatively correlated genes and vice versa. This result reinforces the main critical pathways altered between groups as previously found in the Kegg cluster in Figure 2A. Kegg enrichment analysis of positive correlation for the BL and CL genes are displayed in Figure S4A and S4B clustered by genes. This result indicates a convergence of results by two different methods, reinforcing the enriched pathways.

**Figure S4.**
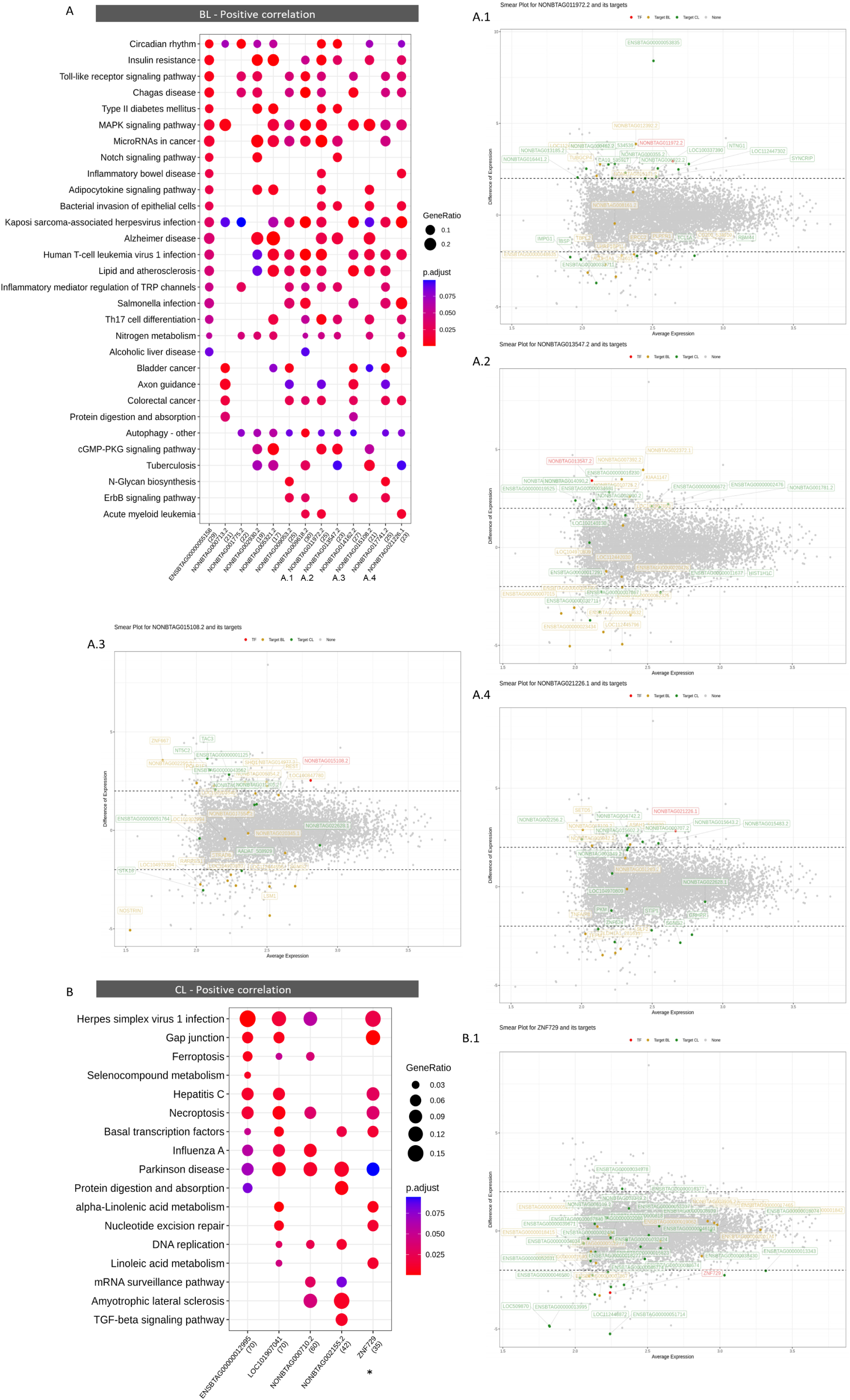
Kegg compare clusters pathways using high correlated genes with overlapped genes. In image A and B we can see pathway enrichment for genes positively co-expressed with each of ovelapped genes. A.1, A.2, A.3, A.4, and B.1 were identified in smearplot analysis using CeTF package showing the gene co-expression network with possible targets. Gray dots represents non-signifficant, red dot is the overlapped gene, yellow and green dots represents respectively BL and CL targets.

We reconstructed gene co-expression network using smearplots from CeTF (Figure S4). In this analysis, we can see the network for each gene based on a different method, the PCIT and RIF. Genes found in the previous analysis appeared in the smearplot co-expression reinforcing their importance. The genes *NONBTAG011972.2* (*TENT5C*), *NONBTAG013547.2* (*DENND11*), *NONBTAG015108.2* (*SEPTIN5*), and *NONBTAG021226.1* (*FOXJ3*) showed a strong correlation of expression with genes that enriched pathways found in the BL group. For example, genes positively correlated with the key gene were associated with RNA stability (*SYNCRIP*, *TENT5C*), nervous system (*NTNG1*), actin cytoskeleton (*ENSBTAG00000053835*), transcription shutdown (*REST, SETD5*), end of maturation (*SHQ1, TAC3, NT5C2, ENSBTAG00000043562 = STK16*) indicating a fine tunning controlled process that is expected in MII oocyte. On the other hand, the genes negatively correlated can be linked to the process of transcription, the beginning of oocyte maturation or oocyte immaturity (*ENSBTAG00000032711 = HTATSF1, IMPG1, IBSP, ENSBTAG00000023434 = BSP1, TEFM*), and DNA damage response (*SLF2, CHKB, STAG2*). In addition, these results corroborates with Molecular Function enrichment.

We constructed a global visualization of the gene network which indicated that *GNAS*, *NFKB1, NRAS, ACTG1, POLRH2, MAPK24*, and *MYC* were the most important genes (in the middle of network with higher connections) for the BL group (Figure S5A). For the CL group, *SLC25A6, PSDM3, PSMC4, TUBB2A, TUBB2B, COX6A1, COX5B, SDHB, LOC104969934, GRIA3, GRIN2A, PSM4, PSMB3* and *BRAF* were the genes participating in a high number of pathways. The Ubiquitin-mediated proteolysis was completely active in the CL group, with a few genes in the BL group. Still, the main pathways were all protein-related, such as Alzheimer, Spinocerebellar ataxia, Prion, Huntington, and Parkinson’s (Figure S5C-E).

**Figure S5.**
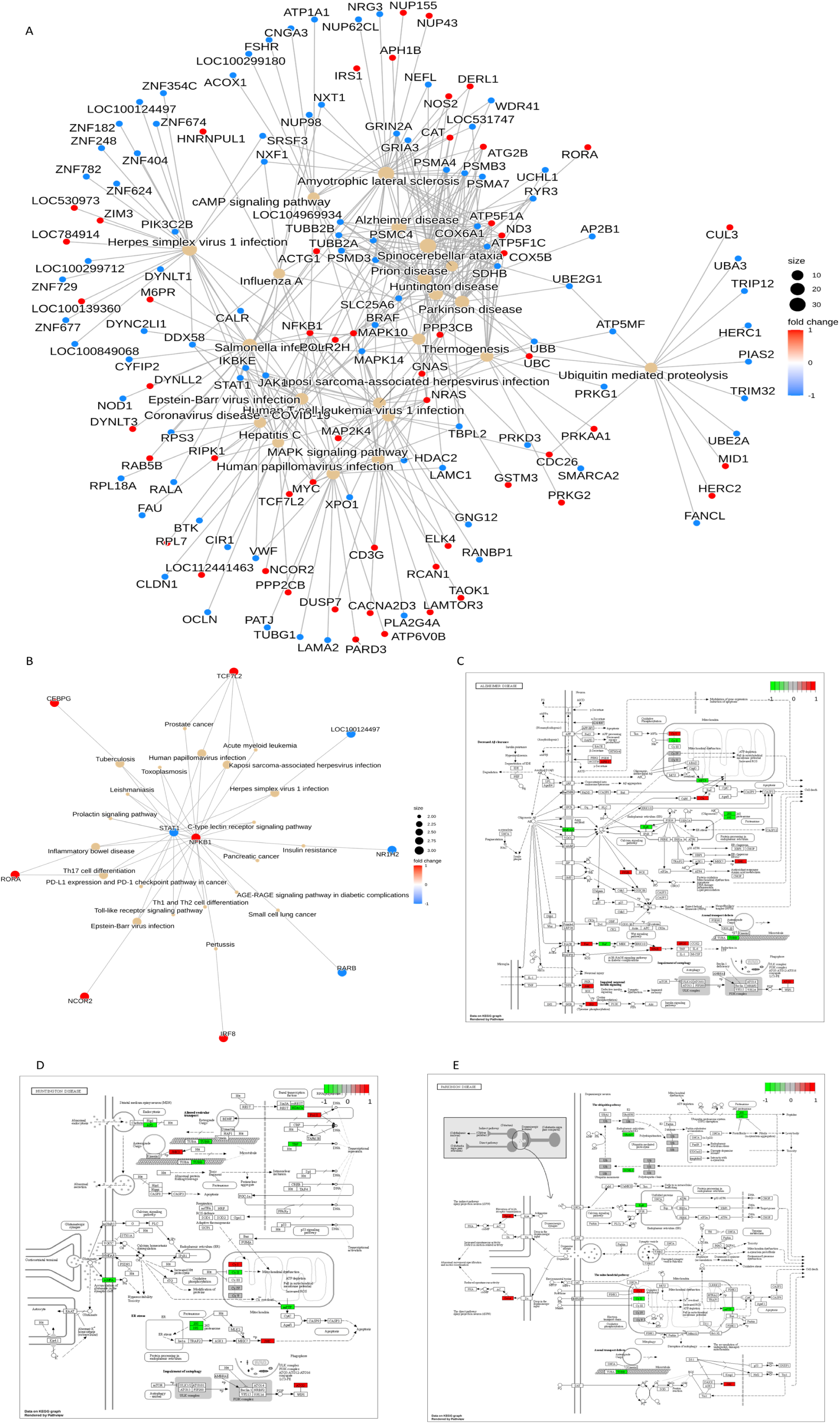
Networks and pathview analysis showing a complete overview of important genes and pathways. A-Network analysis comparing BL and CL groups with the main pathways using merged gene list. B – Network analysis using only non-coding transcripts. Pathview of Alzheimer disease (C), Huntignton disease (D), and Parkinson disease (E) showing molecular mechanisms which are conserved besides the disease name. For pathview, red color represents upregulated gene, while green represents downregulation.

The global visualization of the gene network of lncRNAs highlight *STAT1* (CL) and *NFKB1* (BL) as the key genes (Figure S5B). They were linked to the same pathways, such as Th1 and Th2 cell differentiation, Th17 cell differentiation, AGE-RAGE signalling, Herpes simplex virus 1 infection, and Toll-like receptor signalling. Other genes were also found in this analysis but in the edge of the network. For example, *TCF7L2, CEBPG, RORA, NCOR2*, and *IRF8* were represented in the BL group, while the genes *LOC00124497*, *NR1H2* and *RARB* were represented in the CL group. Among the 580 genes found previously in CeTF analysis, we concatenated gene lists constructed for each RNA classification (lncRNA, protein-coding, pseudogene and TF - Supplementary file 3) and summarised the table in order to count occurences of each one. The genes *GRIN2A* (Glutamate ionotropic receptor *NMDA*), *C7H19orf67* (Chr 19 open reading frame 67), *PIGY* (Protein Pre Y mitochondrial), *PISD* (Phosphatidylserine Decarboxylase Proenzyme mitochondrial), *PRR14L* (Chr 22 open reading frame), *PYURF* (*PIGY* upstream open reading frame) and *SAMD1* (Sterile alpha motif domain containing 1) showed us occurrences higher than 4 among all RNA biotypes identified. These higher occurrences suggest that these genes also can have an important role in oocyte competence affecting a greater number of other genes in both groups.

### *In vitro,* cumulus cells from MII oocytes are not good candidates for investigating oocyte competence

We next investigated cumulus cells transcriptometo investigate the association with oocyte quality. This RNA-Seq was performed in pooled samples comparing BL and CL groups. In our results, we could not find a huge difference in transcriptome analysis between the BL and CL groups (Figure 3).

**Figure 3.**
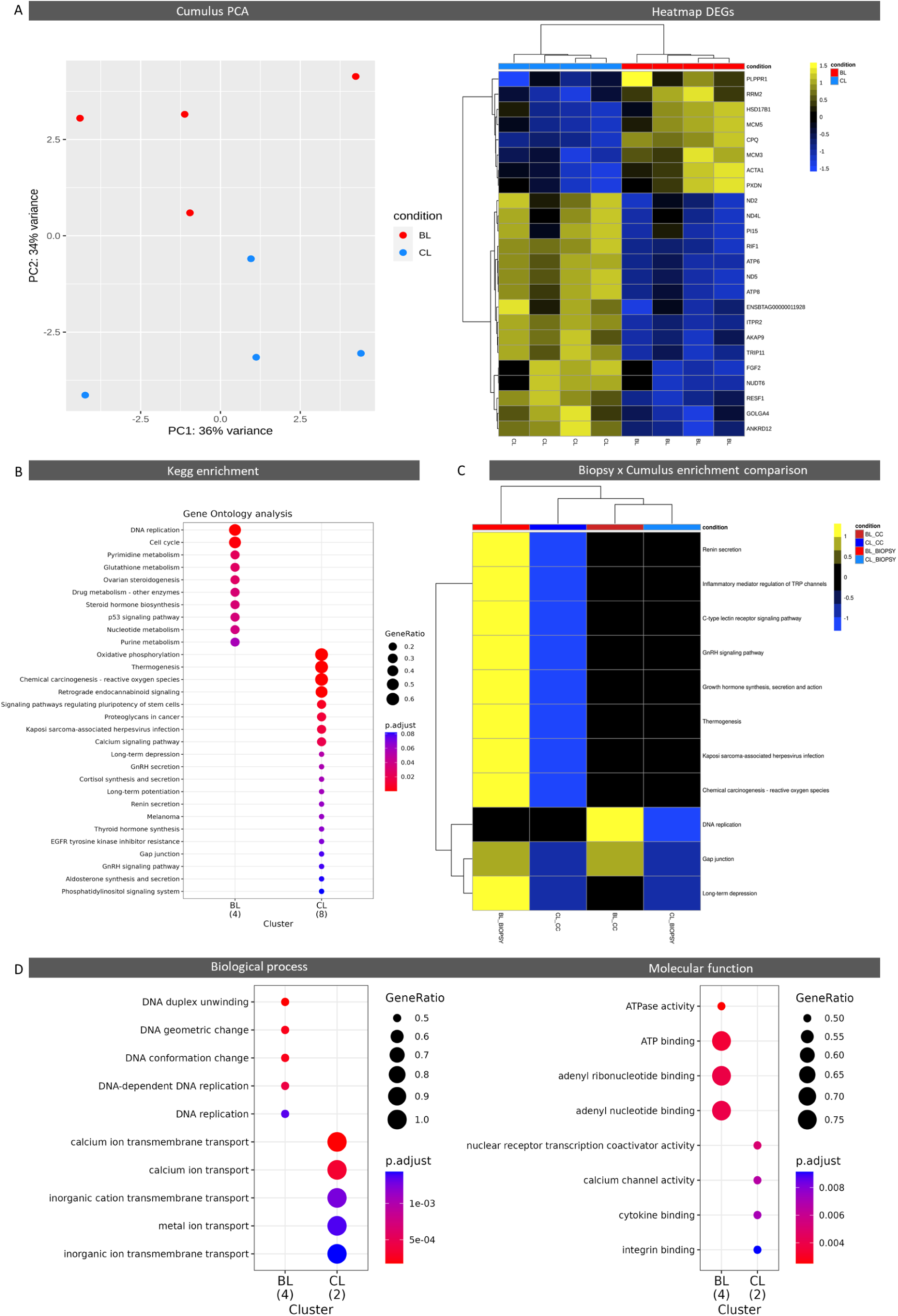
Cumulus cells transcriptome analysis. (A) PCA showing accumulated variance between BL and CL groups and Heatmap containing all DEGs. (B) Kegg compare cluster enrichment using DEGs. (C) Heatmap comparing Kegg enrichment pathways for cumulus and oocyte biopsies. The nomenclature CC and BIOPSY was respectively for cumulus cells and oocyte biopsies. BL group was colored as reds while BL as blues. Active pathways in BL were represented as yellow while blue colours was designated for CL; black colour represent non enrichment; Adjusted p-value was used a continuous color in the palette range where the colour edges contains the smaller adjusted p. (D) Biological proces and molecular function enrichment comparing the cumulus cells transcriptome.

PCA analysis showed a non-complete separation of groups and a poor variance in PC1 (36% of variance) even using the 500 genes with higher variance. We found a total of twenty-four DEGs in this analysis and no exclusive genes. Still, the enrichment analysis indicated that DNA replication, cell cycle, Pyrimidine/Glutathione metabolism, and ovarian steroidogenesis were the main pathways modulated in the BL group (Figure 3B). Moreover, oxidative phosphorylation, thermogenesis, Chemical carcinogenesis - reactive oxygen species, retrograde endocannabinoid signaling, and Signaling pathways regulating pluripotency of stem cells were modulated in the CL group. Additionally we constructed a heatmap using common pathways found in biopsy enrichment and in cumulus enrichment in order to clearly see conserved processes (Figure 3C). This analysis resulted in a clusterisation (from inside to outside) of CL biopsy with BL cumulus group, and CL cumulus with BL biopsy. The cumuls BL group had the biological process pointing to the DNA processes (duplex unwinding, geometric change, conformation, replication), while the CL group was related to calcium and ion transport (metal, transmembrane, inorganic cation transport) (Figure 3D). Molecular function enriched for ATPase activity and binding for the BL group, while cytokin and integrin binding, calcium channel activity and nuclear receptor transcription coactivator activity for the CL group.

### Polar body methylation is associated with developmental competence

The methylation status of the first polar body (1st PB) was investigated using PB from the same oocytes utilized in RNA-Seq. The samples were processed with single-cell WGBS, however we lacked the untreated once the entire sample was bisulfite-treated to determine oocyte genome methylation. Previous studies in humans and mice demonstrated a significant relationship between first PB methylation and the oocyte genome (Yuan et al., 2020; Wei et al., 2019). After peak calling and annotation, we confirmed overlapping methylation genes and regions among samples from the same group, with at least three samples in each group. The distance to transcription sites varied across groups BL and CL, with the majority of the variations happening on the positive strand. The median peak distribution of the CL group was 2000 bp, with a density peak at 1000 bp. The BL group’s median was around 2500 bp (Figure 4).

**Figure 4.**
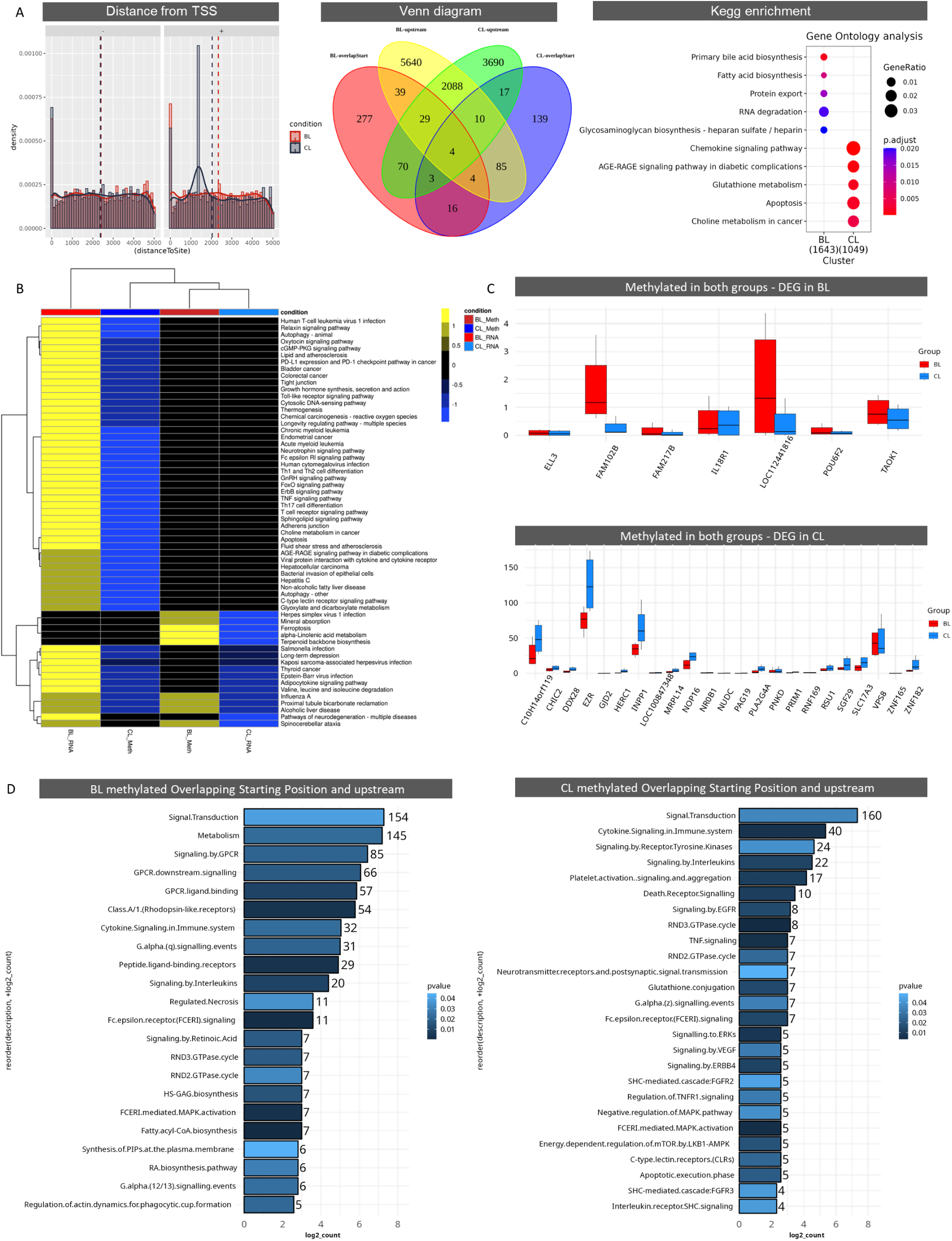
Single WGBS for polar body comparing BL and CL groups. (A) Distance (base pair) from Transcription start site analysis comparing groups and strand showing a few global differences and a changing in median line in positive strand. The Venn diagram resulted in higher differences when genes were compared for “overlapStart” and “upstream”. A total of 5956 and 3846 genes were found respectively for BL and CL groups after MACS3 peak calling. From those genes, the kegg enrichment analysis showed us that these pathways were methylated in one group and not in the other one. (B) The Kegg enriched pathways for methylated sites and for the biopsies RNA-Seq result were merged and than the heatmap comparison was performed. The BL_RNA (RNA- Seq) and BL_meth (methylation) were colored using red and dark red respectively. The CL_RNA and CL_meth were colored using blue and dark blue respectively. Black colour represent non enrichment; Adjusted P value was used a continuous color in the palette range where the colour edges contains the smaller adjusted p. (C) The genes that were methylated and expressed in the same group were plotted in TPM to be investigated as genes which possible were expressed before MII phase. (D) Reactome enrichment was performed using the genes only found in one group and not in the other

The Venn figure illustrates the key different genes, which are subsetted by overlapping start and upstream regions, yielding 5956 and 3846 methylation genes, respectively, for the BL and CL groups. Kegg enrichment was used to identify methylation genes for main bile acid biosynthesis, fatty acid biosynthesis, glycosaminoglycan biosynthesis, protein export, and RNA degradation in the BL group. The CL group, on the other hand, was enriched in chemokine signaling, the AGE-RAGE signaling pathway in diabetic complications, glutathione metabolism, apoptosis, and cancer choline metabolism. Kegg biopsy enrichment and Kegg methylation enrichment were performed to determine the key differences. The heatmap cluster analysis of enrichments indicated that CL_RNA and BL_meth genes were closely related, whereas CL_RNA and BL_meth genes were more comparable. This was in contrast to FoxO signaling, ferroptosis, cGPM-PKG signaling, and GnRH signaling, among other things. Figure 4B demonstrates that PB methylation coincides with gene expression data for the vast majority of pathways that must be shut off at the end of maturation. Despite the fact that the principal peaks were in the differences, several genes were methylated in both groups but expressed differentially in one.

We investigated these genes by limiting them to DEGs. BL DEGs were identified as *ELL3, FAM102B, FAM217B, IL18R1, LOC112441816, POU6F2*, and *TAOK1*. CL DEGs were identified as *RNF169, PRIM1, ZNF182, INPP1, NR0B1, HERC1, VPS8, C10H14orf119, GJD2, DDX28, PNKD, NUDC, ZNF165, CHIC2, NOP16, RSU1, SLC17A3, PAG19, PLA2G4, LOC100847348, MRPL14, EZR, and SGF29*. A series of boxplots were created for each comparison using TPM data to highlight the significant differences (Figure 4C). In addition to Kegg, the Reactome database was used to examine enrichment pathways for methylation genes. Despite differences in gene number, both groups enriched in signal transduction, cytokine signaling in the immune system, and interleukin signaling. The GPCR pathways were different across all populations. GPCR downstream subunits (alpha, beta, and gamma), GPCR subunit-q (related with calcium release), G-alpha (12/13), fatty acyl-CoA

### Granulosa cells transcripts are associated with developmental competence

Following that, we examined the transcriptome of the granulosa samples retrieved from the dissected follicles utilized to acquire the oocytes, as well as their related biopsies, which had previously been categorized as competent (BL) or non-competent (CL). The granulosa samples’ principal component analysis revealed 97% of the variation in the x-axis (Figure 5A). The transcriptome of Granulosa revealed 10287 DEGs (9819 from BL and 468 from CL), 299 unique genes (all from the BL group), and 1424 HubGenes. The heatmap of the fifty genes with the highest variation revealed a strong clustering, with the genes *ENSBTAG00000043567, COX1*, and *ENSBTAG00000043545* (all mitochondrial genes) exhibiting downregulation (Figure 5B). The remaining 47 genes were increased, suggesting dephosphorylation, acetyltransferase, transporter, and axon processes. *LOC101908548, IGF2BP1, LOC104973374, LOC112445622, LOC523769, LOC101902691, PPP1R17, ENSBTAG00000053577, LOC101907485, C13H10orf113, ENSBTAG00000051221* were the genes with the greatest foldChange for the BL group; these genes were associated to mRNA stability, steroid metabolism, and the immune system. *LPAR6, KRT8, TNFRSF12A, LOC101904591, SERPINE1, TIMP1, ENSBTAG00000055072, TWIST1, RGS2, ENSBTAG00000050141, APOA1*; these genes had a closer relationship with transcriptional shutdown, GPCR (G protein-coupled receptor) down-regulation, apoptosis, among the non-coding genes found without description. Furthermore, two ensemble codes (*ENSBTAG00000055072* and *ENSBTAG00000050141*) were labeled to the same microRNA, *bta-mir-2345*.

**Figure 5.**
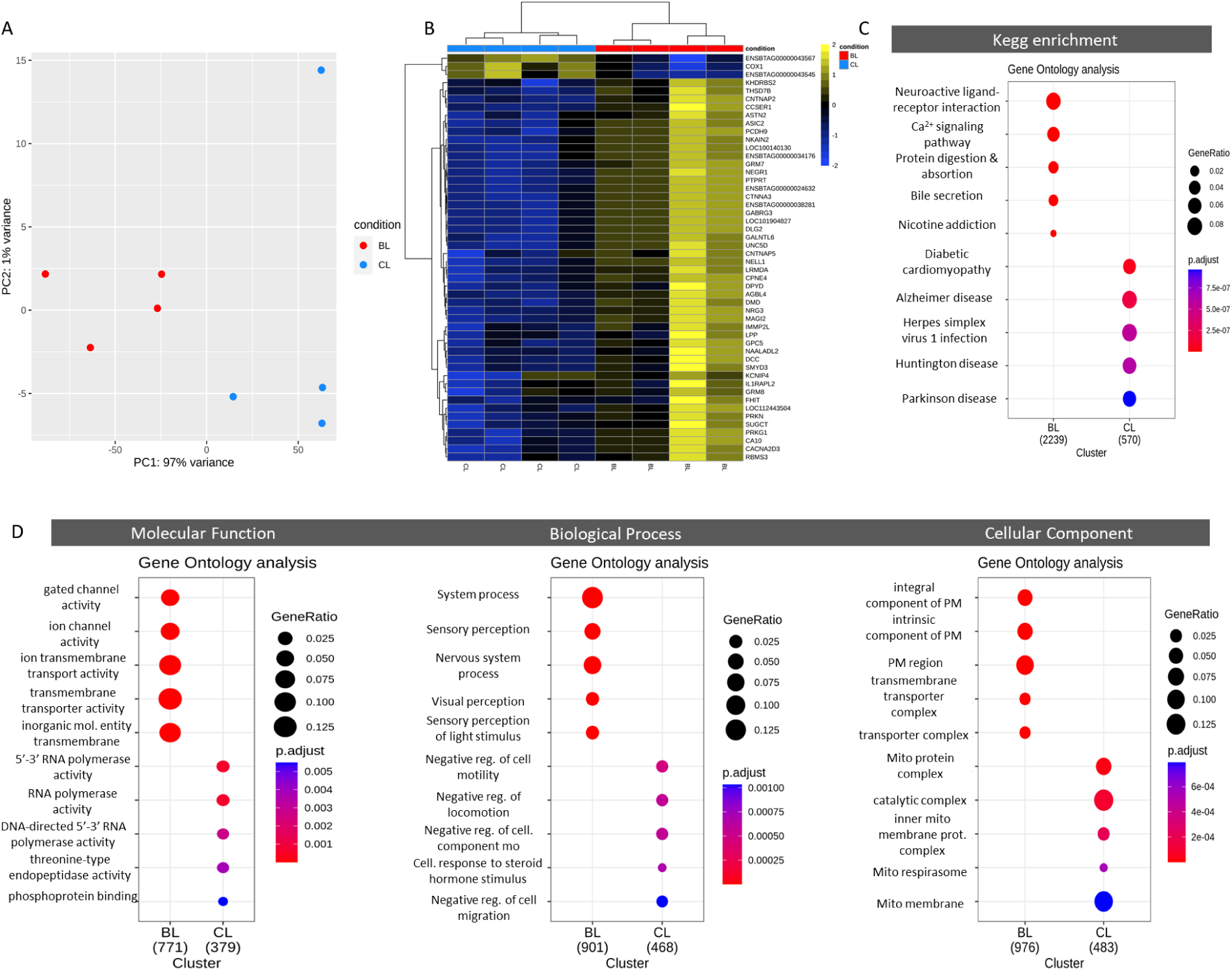
Molecular differences between granulosa transcriptome of BL and CL groups. (A) PCA showing 97% of variance in x-axis. (B) Heatmap of fifty highly variated genes using vst values. (C) Kegg pathways of compare cluster result using DEGs + exclusives + Hubgenes. (D) Gene ontology analysis. They demonstrate the activity that gene product performs (MF), the specific objective that those genes can be programmed to achive in the organism (BP), and where the product can have a function (CC). PM = plasma membrane; Mito = mitochondria

To study the biological processes linked with oocyte competence, we used gene ontology (GO) analysis, reducing redundancy by employing DEGs, Exclusives, and Hubgenes (Figure 5C, Supplementary file 4). Cluster comparisons with Kegg pathways revealed that the BL group contained Neuroactive ligand-receptor interaction, Calcium signaling, Protein digestion and absorption, Bile secretion, and Nicotine addiction. The CL group was enriched for diabetic cardiomyopathy, Alzheimer’s disease, Herpes Simplex Virus 1 infection, Huntington’s disease, and Parkinson’s disease. Furthermore, network analysis for the top five Kegg pathways was performed using a combined gene list (Figure S6).

**Figure S6.**
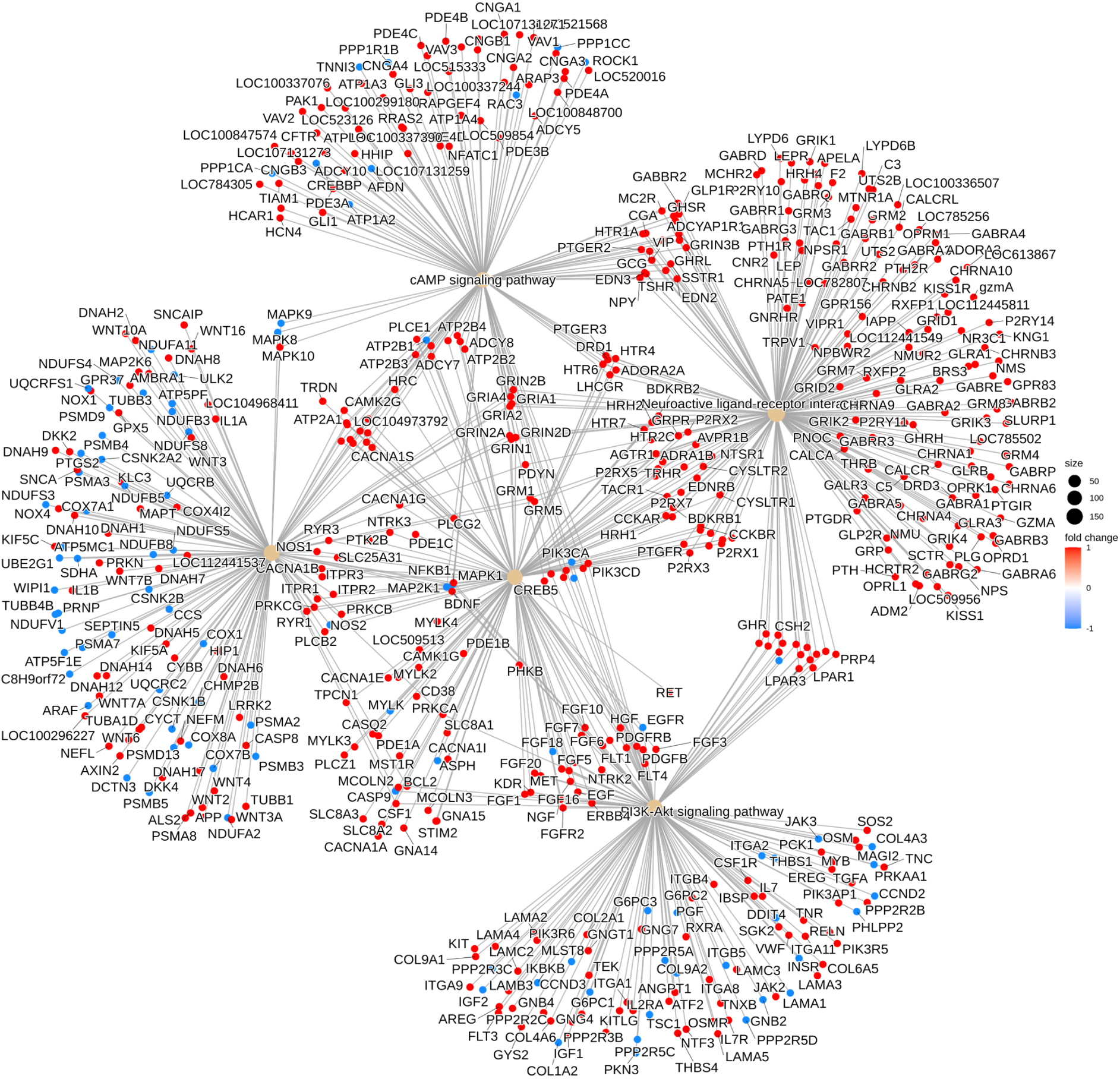
Network analysis of five first pathways. Five most affected pathways in granulosa cells transcriptome. Calcium signaling pathway is the most central pathway, at his left side, the Pathway of Neurodegeneration - multiple disease.

These pathways were very active in the BL group, indicating that the most important genes (higher connections) in the graphic’s center were related to the Calcium and Neuroactive ligand-receptor interaction pathways. The Neurodegeneration-Multiple Diseases pathway, on the other hand, contained the most genes in the CL group. According to gene ontology, the high competence group was involved in the transport of ionic and inorganic chemicals through gated channels and transmembranes. The objective of “feeling” the environment (System perception, Nervous system, Visual perception) and function in the plasma membrane is suggested by the Biological Process analysis. The poor competence group, on the other hand, had activities related to DNA and RNA-polymerase, endopeptidase phosphoprotein binding, indicating a relationship to negative regulation of motility, locomotion, and migration, as well as mitochondrial function (Figure 5D).

### The transcriptome of granulosa cells from competent follicles correlates with in vivo-derived bovine granulosa transcriptome

The gold standard for oocyte quality is cumulus-oocyte complexes exposed to the follicular environment in vivo (Ferré et al., 2020). Because our granulosa cell samples were obtained before the final stages of follicular growth that lead to ovulation, they had not been exposed to the stimulatory effects of an LH peak. Then, we compared them to samples of granulosa data (GSE121588) obtained in vivo before (GnRH0h) and 6h after (GnRH) the LH peak (Schuermann et al., 2018). To eliminate methodological bias, all data were processed following the methods described in the method section. Furthermore, in the following comparisons, we utilized all of the DEGs (10.303) from our BL x CL granulosa transcriptome study. Unsupervised PCA utilizing all genes revealed a PC1 of 77% with the BL and CL groups on opposing sides. With 12% variance, the GnRHs groups were divided by y-axis (Figure 6A).

**Figure 6.**
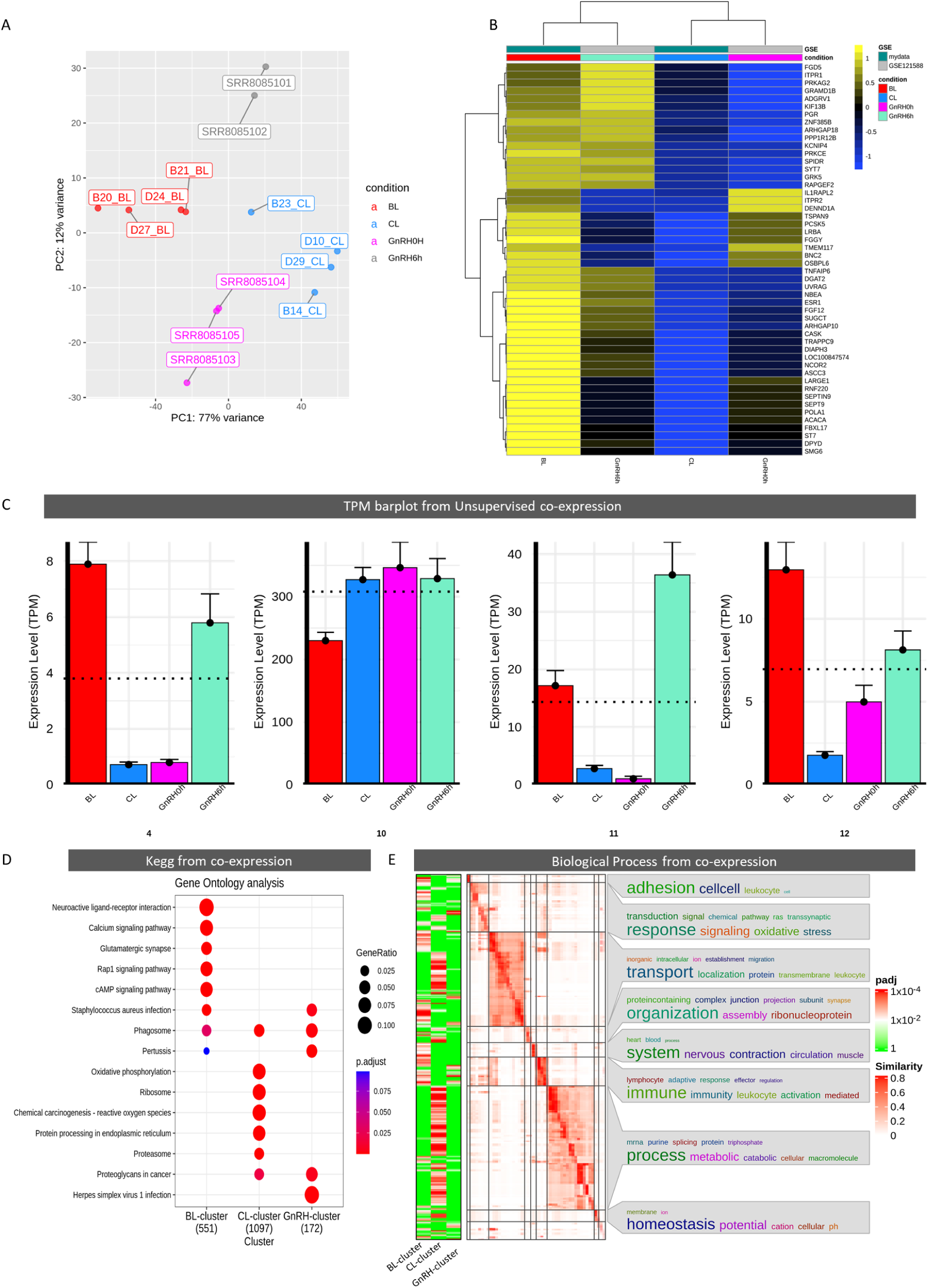
Unsupervised clusterization methods identifies similar clusters. (A) PCA analisis for ou granulosa data and public data. (B) Top fifty genes with highest variance showing common genes with BL and CL groups. (C) Barplot from SOTA algorithm results after identification of common clusters between BL and GnRH groups (4, 11, 12) and between CL and GnRH groups (10) among the thirteen clusters. (D) Kegg enrichment for SOTA and CEMiTool modules were assembled in BL cluster (4, 11, 12, and module 1 from CEMiTool - see supplementary file 5), CL cluster (10), and GnRH cluster (third SOTA cluster). (E) Simplified Biological process from SOTA and CEMiTool modules with the most common terms found in all BPs.

Further analysis with the fifty (50) more expressed genes revealed a somewhat different clustering pattern, with the BL group remaining comparable to the GnRH6h group and the CL group clustering with the GnRH0h group (Figure 6B). *FGD5, ITPPR1, PRKAG2, GRAMD1B, ADGRV1, KIF13B, PGR, ZNF385B, ARHGAP18, PPP1R12B, KCNIP4, PRKCE, SPIDR, SYT7, GRK5*, and *RAPGEF2* are among the highest expressed genes in both BL and GnRH6h. The genes shared by BL and GnRH0h included *IL1RAPL2, ITPR2, DENND1A, TSPAN9, PCSK5, LRBA, FGGY, TMEM117, BNC2, OSBPL6, TNFAIP6, DGAT2*, *UVRAG, NBEA, ESR1, FGF12, SUGCT, ARHGAP10, CASK, LOC100847574, NCOR2,* and *ASCC3*.

To identify potential granulosa biomarkers associated to oocyte competency, an unsupervised analysis was done using SOTA (Self-organizing tree method) and CEMiTool. Twelve (12) clusters were discovered for SOTA. Clusters eleven (11) and twelve (12) exhibited greater BL and GnRH6h co-expression. Cluster ten (10) displayed co-expression between the CL and GnRHs groups, whereas module three had solely GnRHs co-expression. CEMiTool discovered three (03) clusters, but only one had a convincing pattern confirming co-expression with BL and GnRHs (Figure S7). The clusters were divided into three types: the BL-cluster (SOTA: 4, 11, and 12; 1710 genes), the CL-cluster (SOTA: 3; 2052 genes), and the GnRH-cluster (SOTA: 10; 723 genes) (Figure 6C). According to KEGG study, the BL- cluster was associated with neuroactive-ligand, calcium, glutaminergic synapse, Rap1 and cAMP, whereas the CL-cluster was associated with the proteasome, protein processing in the endoplasmic reticulum, proteoglycans in cancer, and Herpes simplex virus 1 infection. The GnRH cluster enriched for staphylococcus aureus infection, phagosome, and pertussis, all of which were also present in the BL-cluster. Cancer pathway proteoglycans shared CL and GnRH clusters (Figure 6D). In co-expression analysis, a simplified picture of the biological process revealed that BL and GnRH clusters were more comparable than CL clusters (Figure 6E). Cell adhesion, response signaling, immune system - leukocyte activation and lymphocyte, inorganic homeostasis, and cation were the most commonly used phrases. The CC indicated a function with an inherent plasma membrane component, such as a projection membrane protein complex, axon, and plasma membrane with molecular functions in gate channel, transmembrane transport, and nucleic acid binding (data not given).

**Figure S7.**
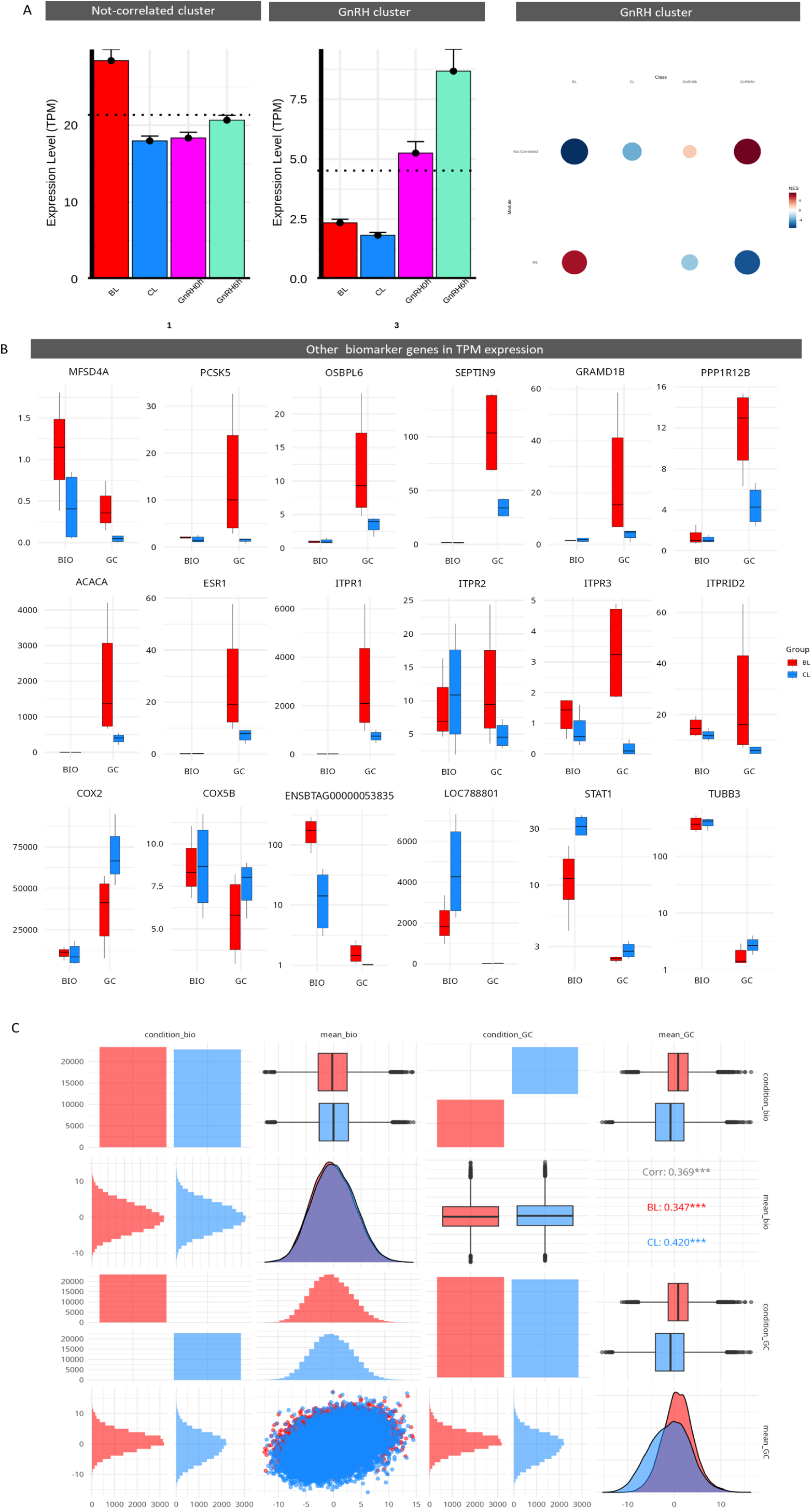
Unsupervised clusters and genes. (A) Cluster from unsupervised analysis showing genes with no signifficant co-expression, GnRH co-expressed genes, and a cluster with BL and GnRHs co-expression. (B) TPM gene expression of biomarkers associated with oocyte quality.(C) A pairwise matrix of granulosa cells, oocyte biopsy, BL, and CL groups showing all differences and similarities. This shows that main difference came from granulosa cells in gene expression levels. The distribution found in oocyte was quite similar between groups. In the dispersion plot we can see a higher expression and number of genes in BL group, but that this came majoritary from BL granulosa cells once CL oocyte showed more DEGs and higher level of expression.

This allowed us to identify potential granulosa biomarkers linked with oocyte competence that were consistent with in vivo data while also being connected with our high competent group. These indicators go beyond established oocyte quality biomarkers, once we did not find any difference in those biomarkers whe BL and CL group were compared (Figure 7A). The parameters TPM counts, log2FoldChange, and Gini stability were used to sort all of these genes and additional genes filtered using a systematic technique (Figure 7B, Figure S7B). As high-quality indicators, the novel biomarker candidates were *FSTL4, IGF2BP1, AKAP6, PCKS2, PRKCE, ADGRV1, DENND1A*, and *TSPAN9*. Low-quality markers were discovered as *COX1, TMSB4X, ATP5F1C, SLC25A6, TUBB2A, EZR, TUG1*, and *FTL*. Such oocyte and granulosa indicators clearly suggest a possible involvement in development fine-tuning, since they are all stable between groups and specific for oocyte (BIO) or granulosa cells (GC). Figure S7B shows the identification and description of other key genes in our study. The dispersion map of the pairwise matrix comparing granulosa, oocyte biopsy, BL, and CL revealed that the greatest differences across groups were from granulosa BL. Furthermore, the CL group had higher levels of expression in oocytes. This might imply that the primary variations occurred in follicles prior to oocyte harvest (Figure S7C).

**Figure 7.**
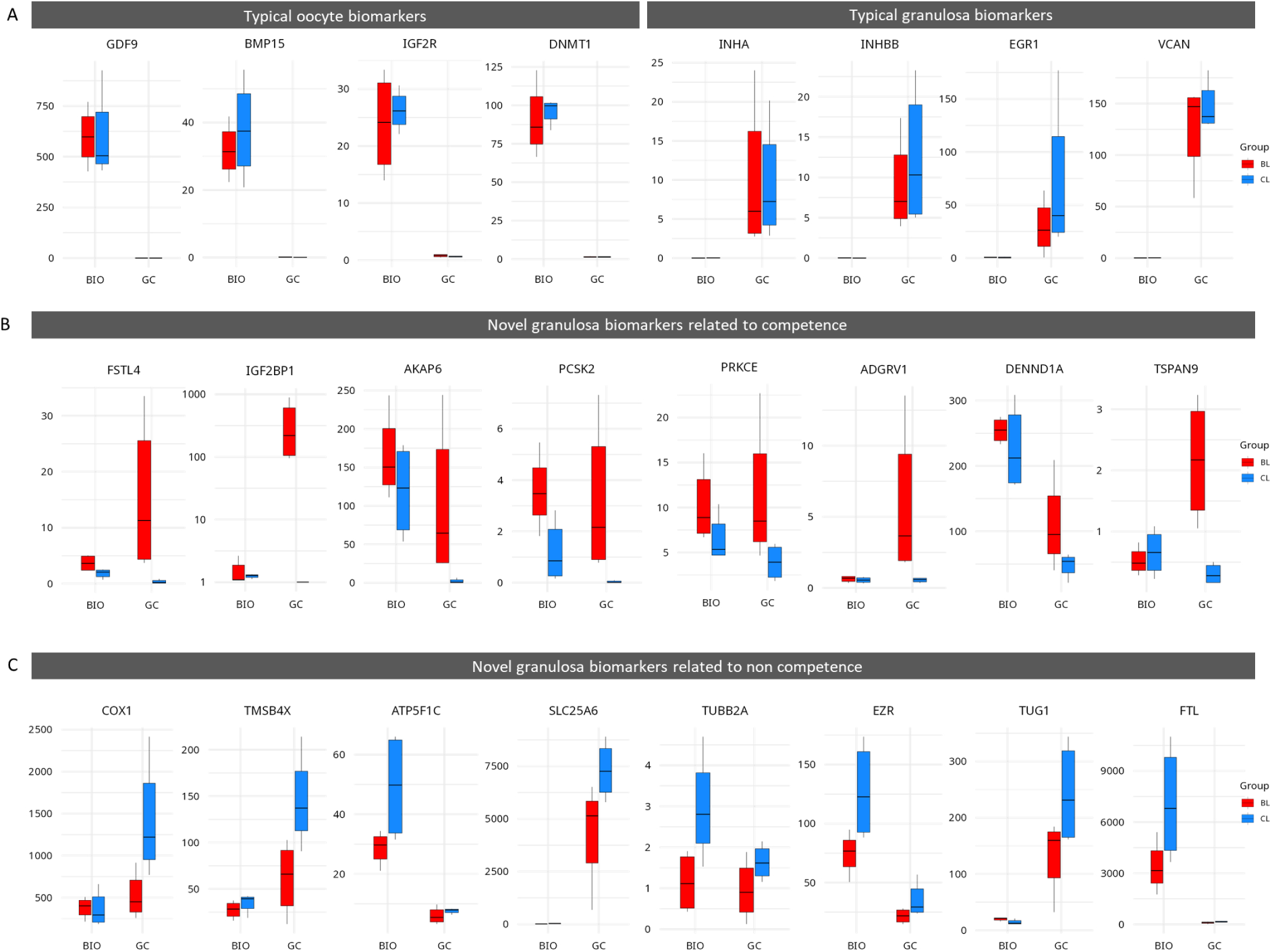
Granulosa biomarkers related to oocyte quality co-expressed with *in vivo* granulosa and sorter by TPM and log2FoldChange. (A) Typical oocyte and grnaulosa biomarkers. (B) Novel granulosa biomarkers related to granulosa folliculogenesis and oocyte competence. (C) Novel granulosa biomarkers related to granulosa folliculogenesis and oocyte incompetence.

## DISCUSSION

Many research have advocated analyzing cumulus cells to better understand oocyte contents because to their close contact and known interactions (Biase & Kimble, 2018). In the past, a similar study investigated oocyte biopsy using microarray approach were they found just a few differences and with the gene *IGF2R* upregulated in a high competence group (Biase et al., 2012). Here, we deeply investigated oocyte RNAs content using a Sc-oocyte-biopsy RNA-Seq approach to characterize the RNAs, pathways, and hub genes found in the MII oocyte. Additionally, we investigated the relationship of oocyte competence with its follicular niche utilizing transcriptome analysis from granulosa and cumulus cells, as well as DNA methylation using the first PB. The approach using single transcriptome and methylome revealed a high number of differences in genes, pathways and methylated regions between BL and CL groups. However we did not find differences in *IGF2R* expression or methylation. This can be due to two main factors, the Single-cell approach used here and the strict follicle size, ∼5mm.

In the BL x CL comparison, our findings revealed a strong phenotypic difference between transcriptomes from biopsies and granulosa cells. The similar thing was observed in the WGBS study. However, just a few differentially expressed genes (DEGs) were discovered in cumulus cells, and they did not cluster completely in PCA. Nonetheless, the enriched pathways discovered in cumulus were completely contrary to those discovered in biopsy analysis and granulosa. The observed disparities in granulosa and oocyte groups imply that they are from different follicular settings. As a result, after subjecting these cumulus to a standardized culture environment, a compensatory gene expression changing was induced, thereby mitigating the differences and masking the molecules associated with the competent and non competent groups. This could mean that the *in vitro* environment is forcing a normalization of gene expression, pushing the non-competent oocyte in the direction of competent (cleavage), but not strong enought. The large differences found in biopsy, which are consistent with granulosa cells, suggests that oocyte transcription is weak or absent during the end of maturation. As a result, RNA preservation occurred prior to in vitro maturation. This theory is backed by the WGBS finding, which found that methylation was tightly associated to gene transcription shutdown.

Oocytes accumulate a variety of transcripts, proteins, and other molecules necessary to support development until maternal-zygotic transition. Our results indicate that the transcripts present in the oocyte and their respective granulosa cells of competent and non-competent groups are distinct and relevant genes, as indicated in heatmap showing the top fifty highly variable genes. These genes turned-out repeatedly in different analytical approaches, e.g. pathways, smearplots, unsupervised clusterisation, and classificatory method, and still, in WGBS analysis suggesting their consistency and importance as potential biomarkers of oocyte quality.

*PCSK2* (proprotein convertase subtilisin/kexin type 2) was found to be overexpressed in biopsies and granulosa cells. This protein is involved in the conversion of precursor proteins into physiologically active peptides that are expressed in a variety of organs, including the ovary, where it has been demonstrated to have a role in follicular development and oocyte maturation. Reduced expression of PCSK5 in granulosa cells may result in suboptimal oocyte maturation and embryo development (Antenos et al., 2011). Furthermore, we discovered the paralogous *PCSK5* as a granulosa DEG and hub gene in the oocyte.

The TGF superfamily is made up of a vast number of proteins and ligands morphogens that are involved in cell proliferation and differentiation (Wijayarathna & de Kretser, 2016). Follistatin, a TGF family ligand, for example, was shown to be positively linked with mRNA levels in competent oocytes from cows (Patel et al., 2007). Furthermore, functional testing in in vitro maturation with *FSTL* ablation revealed a consistent impact in which it enhanced the quantity of cow blastocysts and trophectoderm cells numbers. The same group discovered no link between follistatin and the GDF9 or BMP15 molecules, suggesting a distinct functional role for follistatin other than activin binding (Kim et al., 2011). We discovered a similar finding with the *FSTL4* gene as one of the most variable genes in the heatmap for the BL group. We also did no observe any difference in *GD9* and *BMP15* levels between the BL and CL groups. Still, we found other TGF-Beta member among heatmap genes in the BL group, SNX25. The sortin nexin 25 counteracts the TGF-Beta pathway by degrading receptors via the lysosomal route (Hao et al., 2011). Interestingly, we discovered the TGF-Beta genes *NEO*, *ZFYVE16* (biopsy hubgenes), *TGIF1*, and *HJV* (DEGs) in the CL but not in the BL groups, indicating that this pathway is active exclusively in the CL group. These findings imply that in MII phase oocytes, the TGF-Beta pathway is being shuting off in the BL group where SNX25 participates on it. Reinforcing that, we found TGF-Beta enriched into the CL group in oocyte.

Aside from signaling pathways, mRNA stability was discovered to be a crucial component in oocyte quality. IGF2BP1, a member of the insulin-like growth factor 2 mRNA-binding protein family, was one of the genes with the greatest log2FoldChange in granulosa. This protein regulates the translation of *IGF2*, beta-actin, and beta-transducin repeat-containing protein mRNAs. Weidensdorfer and coworkers discovered that IGF2BP1, SYNCRIP, HNRNPU, YBX1, and DHX9 bind to the Coding Region of the MYC RNA and help to stabilize it (Weidensdorfer et al., 2008). Despite the fact that *IGF2BP1* was only found in granulosa cells in our investigation, it was deemed a useful biomarker for competence. Furthermore, *SYNCRIP* was one of the fifty genes with the largest variance that were elevated in the oocyte. We discovered *MYC* expression increased in biopsy with a log2FoldChange of 6.76, comparable to Weidensdorfer, which may imply a similar function for IGF2BP1 and SYNCRIP as key genes to maintain MYC, an important transcription factor. Furthermore, *NONBTAG011972.2* (*TENT5*), a hub gene and DEG in oocytes, was one of the fifty highly expressed genes in the oocyte’s heatmap. *TENT5* function is also connected to mRNA stability during gene expression via elongation ending with the uridine molecule, which supports the previous findings (Mroczek et al., 2017). The coexpression patterns for this noncoding RNA were enriched for cGMP-PKG, autophagy, axon guidance, and MAPK signaling, which may be dependent on mRNA stabilization factors (Figure S4A, 4A.1).

Phosphorylation is one of the most common methods of signaling in most cells (Ardito et al., 2017). PRKCs are a kind of protein kinase that may be triggered by calcium and the second messenger diacylglycerol. Calcium was shown to be one of the most significant secondary messengers in cells and one of the top five most regulated pathways in granulosa cells from the BL group (Stewart & Davis, 2019). In mice, maternal PRKCE buildup in mature oocytes is required for the first cleavage, which facilitates the maternal-to-zygotic transition (Zhang et al., 2022). This gene was discovered to be increased in granulosa cells and oocytes, indicating an essential function and potential early biomarker candidate because it may be identified before maturation. Another kinase-related protein, *AKAP6* (A-kinase anchor proteins), which regulates cAMP pathway by binding to the regulatory subunit of protein kinase A and confining it to a specific area of the cell (Dodge-Kafka et al., 2005). Moreover, *AKAP6* is implicated in oocyte quality and meiosis, it has been linked to the regulation of oocyte mitochondrial function, and was shown to be downregulated in human oocytes from old women (Brown et al., 2002; Vergarajauregui et al., 2020; Zhang et al., 2020). We showed that AKAP6 was widely expressed and upregulated in both, in granulosa cells and oocytes competent groups. Furthermore, the cAMP pathways were modulated in BL granulosa while oocyte meiosis in oocytes. In addition, other *AKAPs* such as *AKAP4, AKAP3, AKAP12*, and *AKAP14* were also found upregulated in granulosa. Other genes associated with this kinase, such as *PRKG1* and *PRKG2*, were shown to be differently expressed in granulosa cells and oocytes, reinforcing the relevance of the cAMP and cGMP-PKG pathways, as well as the *AKAP6* gene as a potent biomarker.

G-proteins are a protein family that is engaged in external to internal signaling transmission and may activate or deactivate cellular processes in response to subunit stimulation (Liccardo et al., 2022). Here, we found the *ADGRV1* (*GPR98*), a member of the G-protein coupled receptor superfamily as an important biomarker upregulated in granulosa cells. This protein is essential in the neurological system, where it binds to calcium to triggers his functions. It couples to G-alpha(i)-proteins and *GNAS*, reducing adenylate cyclase activity and cAMP production. *GNAS* (also known as *GNAS* complex locus - G protein alpha subunit group S) is an maternally expressed imprinted locus that can produce different transcripts depending on methylation status and has been identified as a key gene for oocyte competence in mice (Li et al., 2020). Ion channels such as atrial voltage gated sodium channels and dihydropyridine-sensitive calcium channels are likewise activated by G alpha-S. These pathways were extremely enriched in granulosa cells and in oocyte. In our study, *GNAS* was classified as a Hub gene, and the two ensembl codes that we discovered here were classified as Hub genes in the BL group (*ENSBTAG00000047223*, *ENSBTAG00000052413*).

Contrast with this result, the G-protein that has the opposite effect of *GNAS*, protein G subunit I (*GNAI1* - adenylyl cyclase inhibition), was shown to be downregulated in the oocyte. Furthermore, the adenylyl cyclases *ADCY5, ADCY1, ADCY7, ADCY8, ADCY10*, and *ADCYAP1R1* (adenylyl cyclase controlled by LH) were all elevated in granulosa cells, suggesting stimulation of pathway prior to oocyte maturation. Reinforcing this results, the WGBS analysis indicated that the GPCRs pathways (Reactome) were enriched in the BL group. Nonetheless, these pathways were mostly linked to positive signaling transduction events. This might imply a fine-tuning coordination of several pathways, as well as a critical gene family that are expressed before or during oocyte maturation and then shutted-down.

Another gene controlled by methylation levels, i.e. *DENND1A* (DENN Domain Containing 1A), is hypermethyated oocytes from old cows, suggesting that low expression of *DENND1A* is associated with advanced maternal age and could consequently be related to poor oocyte quality (Liu et al., 2021). Here, we found DENND1A upregulated in granulosa and also other genes from the same family: *DENND2C, DENND2D, DENND5B, DENND2A, DENND11* (*NONBTAG013547.2* - DEG and hub gene); *DENND1A* and *DENND1B* were also upregulated in granulosa cells.

In a human oocyte study, *TSPAN9*, an estrogen responsive gene, was shown to be downregulated in blastocyst from old women and was associated with poor development (ankovičová et al., 2020). We showed that *ESR1* was upregulated in granulosa and the genes *NONBTAG014862.1* and *NONBTAG014862.2* identified in heatmap analysis (both blasted as *SMIM14*) have been shown to be important for blastocyst hatching. while the gene *ENSBTAG00000053835* has been identified in bovine cumulus as a biomarker of oocyte quality (Walker & Biase, 2018 - SSR Abstract).

In the co-expressoin analysis for genes foun as DEGs and Hubgenes in non competent CL group, the genes *LOC101907041*, *MCUB*, *ZNF729*, *NONBTAG002155.2*, and *NONBTAG000710.2* were correlated (corr > 0.75; p-value 0.05) with genes enriched for the most important pathways found in the CL group, such as necroptosis, ubiquitin-mediated proteolysis, gap junction, TGF-Beta, and protein digestion and absorption. *ZNF729* is a DNA-binding transcription factor that positively regulates gene transcription. The smearplot co-expression network (Figure S4) indicated that together with this gene, the genes positively co-expressed were also involved in a positive regulation of transcription, indicating a molecular immaturity of non-competent oocyte oocyte. This result is reinforced by the cumulus cells, once we speculate that the aberrante expression found in cumulus can be a result of an artificial environment induction.

Corroborating with the fact aforementioned that MII oocyte must be prepared to support development until maternal-zygote transition, the first biomarker associated with poor embryo quality was de *COX1*, a component of mitochondrial cytochrome C oxydase which is the component of the respiratory chain that catalyses the reduction of oxygen to water. The number of mitochondria increases following folliculogenesis (Kirillova et al., 2021). However, when oocytes achieved full maturation, it displays low mitochondrial activity (KOYAMA et al., 2014). The high expression of *COX1* represents an oxidative stress process. In porcine oocytes, the melatonin treatment could improve oocyte maturation and embryo development with a significant decrease of *COX1* mRNA and protein levels. At the same time, it could increase *COX5B*, another component of cytochrome C (Niu et a., 2019; He et al., 2017). In our findings, *COX1* was shown to be downregulated in granulosa cells and oocytes in our study. Reinforcing melatonin results, *COX5B* was identified as an oocyte Hub gene linked to the competent BL group.

Indirectly related to mitochondrial activity, another CL biomarker was *TMSB4X*, which codifies a Thymosin Beta 4 X-Linked that negatively regulates actin polymerization thought sequestration of actin G (Zhu et al., 2019, Skruber et al., 2018). Other *TMSBs* genes were found in granulosa and oocyte transcriptome data, where the *TMSB4X, TMSB10*, and *TMSB15B* were also found downregulated in granulosa. Despite this, all of them were largely expressed in oocytes and in granulosa, suggesting a canonical role. *Tmsb4x* was found to be upregulated in mouse granulosa after overexpression of *Slc25a26*, which may have triggered a decline in oocyte maturation by causing oxydative stress (Cheng et al., 2022).

Yang and colleagues (2022) found that *Atp5f1c* (ATP synthase F1 subunit gamma) is critical for mouse oocyte maturation. It was shown to be significantly expressed in granulosa cells and oocytes (6.80 and 39 TPM, respectively), but downregulated exclusively in the oocyte, with a propensity to be downregulated in granulosa cells. In oocytes, the mitochondrial gene *SLC25A6* was likewise downregulated. This gene belongs to the subfamily of solute carrier proteins known as mitochondrial carrier genes. The product of this gene functions as a gated pore that translocates ADP from the cytoplasm into the mitochondrial matrix and ATP from the mitochondrial matrix into the cytoplasm. In cat oocyte it is largely expressed in immature oocytes while this protein appears to be negatively correlated with sperm fertilizing ability in humans (Lee et al., 2017; Torra-Massana et al., 2021). Therefore, this corroborates with the idea of a molecular delayed oocyte in CL group.

Tubulin are globular proteins that polymerize into microtubules, which are part of the cytoskeleton in eukaryotic cells, where it participates in meiosis, mitosis, and mitochondrial distribution. During meiosis and mitosis, there is a high energetic demand for tubulin in specific regions of the oocyte, indicating it plays an important role (Matthews et al., 1993). Here, we found *TUBB2A* (Tubulin Beta 2A Class IIa) highly expressed in both granulosa cells and oocytes. However, *TUBB2A* downregulation in granulosa and oocytes suggests it is required for high mitochondrial activity and/or transport, which is characteristic of a non MII oocyte. In addition, *TUBB2B* was also found downregulated in granulosa and in oocytes, as *TUBB3*. Together, the mitochondrial related genes with granulosa / oocyte Cellular Component enrichment (mitochondrial pathways and high energetic dependent pathways egg: spindle) converged to an idea that non competent oocytes and their associated cells in the follicle are immature at the molecular level, as aforementioned, since mature oocytes displays low mitochondrial activity (Van Blerkom, 2011). Still, in oocyte we can see pathways reinforcing our thoughts such as, DNA replication, TGF-Beta, Nucleocytoplasmic transport. Reinforcing this, we found the gene *TAOK1* (TAO kinase 1) as a DEG, and at the same time methylated in the BL group suggesting that his expression occurred possible during or before maturation and must be repressed at the end of oocyte maturation. This gene triggers microtubules instability and disassembly in neuronal studies (Beeman et al., 2023).

Together, the mitochondrial related genes with granulosa/oocyte Cellular Component enrichment (mitochondrial pathways and high energetic dependent pathways egg: spindle) converged to the idea that this non competent oocyte and associated cells in follicle are molecularly delayed, as previously stated, and mature oocytes have low mitochondrial activity (Beeman et al., 2023). Still, in the oocyte, we may find pathways that support our ideas, such as DNA replication, TGF-Beta, and nucleocytoplasmic transport.

We also showed that *EZR* (ezrin) was downregulated in oocytes and to have a propensity to be downregulated in granulosa (p-value 0.06, padj 0.15). It is a cytoplasmic peripheral membrane protein that functions as a link between the plasma membrane and the actin cytoskeleton. This might be connected to the tubulin changes discovered in the CL group. Furthermore, the downregulation of this gene in bovine oocytes thought micro-RNA bta-mir-183 could substantially improve cleavage and blastocyst development (Wu et al., 2023), suggesting that, despite of its importance in cytoskeleton and cell integrity, overexpression may be associated with poor embryo development.

Aside from mitochondrial-related transcripts, we found that lncRNA *TUG1* (Taurine upregulated 1) expression was associated with incompetence. Long-noncoding transcripts make up a considerable portion of the genomes of several species, and are usually species specific (Noviello et al., 2018). As an exception, *TUG1*, Taurine upregulated 1, is an example, since it is an evolutionarily highly conserved lncRNA that was first recognized as essential in retinal development of rodents (Young et al., 2005). Since then, TUG1 has been implicated in diverse biological processes, including chromatin remodeling, or acting as a trap for proteins or miRNAs. Interestingly it was one gene downregulated in granulosa cells but upregulated in oocytes. In addition, *MEG3*, another important maternally expressed lncRNA, was found upregulated in granulosa and with a tendency to be upregulated in oocyte (3.35 log2FoldChange, padj 0.11).

Ferroptosis was found downregulated in non-competent oocytes. The genes that participate in this pathway, *SLC40A1* and FTL, were downregulated in both oocytes and granulosa cells, but *FTH1* and *LOC788801* were downregulated solely in granulosa cells. In general, iron overload has been linked to infertility and oocyte dysmaturity in humans. It also risks of endometriosis (Ni et al., 2022). Interestingly, the ferroptosis pathway was modulated in BL granulosa cells while it was modulated in CL group in oocytes, suggesting the initiation of a cellular death program in granulosa and in oocytes, which may be reinforced the necroptosis pathway as identified as activated in the oocyte CL group.

Furthermore, variations between the BL and CL groups were clearly connected with optimal maturation in biopsies and granulosa cells, but not in matured cumulus cells. Most pathways in the BL group had previously been linked to physiological oocyte/follicle development, including MAPK, FoxO, GnRH signaling, cGMP, TGF-Beta, Oocyte meiosis, and Erbb signaling. Once MAPK and FoxO, for example, are the key pathways contributing to normal meiosis progression (Sun et al., 1999; Kuscu & Celik-Ozenci, 2015). These pathways represent together the true oocyte requirement for development to the blastocyst stage, once MAPK and FoxO for example are the main pathway contributing to correct meiosis progression. Furthermore, GnRH pathway was present in granulosa and oocytes from the competent BL group. This indicate that these granulosa cells were responsive to the LH hormone, which is directly associated with the cGMP pathway in granulosa, *ERK1/2* and AREG/EREG in cumulus (found upregulated in granulosa BL), and oocyte factors, such as GDF9 and BMP15 (Nunes et al., 2015). Despite this, there was no change in *GDF9*, *BMP15*, or other oocyte factors when comparing BL and CL groups. Interestingly, the.

Neuroactive ligand pathway, is an important communication network between ocytes and its surrounding cells (Biase & Kimble, 2018). We found this pathway hyperstimulated in granulosa cells from the BL group. This pathways is associated with ovarian follicle growth, differentiation follicle secretion and maturation (Sun et al., 2021). In our study, all genes in this pathway were activated in the BL group and notably, among receptors in this pathway, *MCHR, FSH, LHB*, *LHCGR, TSH*, and *TSHR* are modulated in the BL group, suggesting a modulatory role by these hormones. Only the S2R (Progesterone Sigma receptor) was downregulated granulosa, despite all oocytes were collected from ovaries with corpus luteum grade III. Reinforcing this, Matoba and collegues used a retrospective study to investigate the metabolomic profile of follicles following criteria similar to the ones utilized in our study (BL x CL). They identified no differences in oestrogen, testosterone, and progesterone levels between these groups in follicular fluid, but differences in receptor mRNA levels in granulosa (Matoba et al., 2014).

## Conclusion

In conclusion, using a single-cell approach to generate a single ooplasm RNA-Seq from a small portion of an oocyte, we were able to retrospectively examine the competence of oocytes and distinguish between granulosa cells and oocyte cells. In addition, the differences in methylation utilizing PB WGBS corroborates with transcriptome analysis being possible another biomarker source, and with no effect in the progenie (Alteri et al., 2023). Our research sheds light on the importance of maternal transcripts in early zygote development and emphasizes the possibility of using granulosa cells from recover during follicular aspiration as early biomarkers of early developmental competence. In addition, our findings elucidate the molecular mechanisms underlying competence, highlighting the essential pathways for blastocyst development that must be completed prior to occyte activation (MII). The enrichment for transcripts in the granulosa of the non-competent CL group indicated a relationship to a non-responsive granulosa following the shutdown of transcription. In contrast, the pathways and genes identified in as biomarkers in the non-competent oocytes indicate a molecular retardation and death fate. Still, our data suggests that oocyte fate is inherited from distinct follicular niches (granulosa transcriptome), which itself is a major influencing factor in oocyte quality while the cumulus cell present after *in vitro* maturation can be a confounding factor due to their compensatory effect caused by the response to the same *in vitro* environment.

## MATERIAL AND METHODS

### Source of oocytes and granulosa cells

The methods used to recover oocytes and granulosa cells (GC) have been described previously (Andrade et al., 2017). Briefly, bovine ovaries obtained from a local slaughterhouses were transported to the laboratory in thermic bottles in sterile saline solution (0.9% NaCl) at approximately 26 °C within an interval up to 3 h after slaughter. Ovaries from eight routines were used to dissect individual follicles with diameters ranging from 3 to 6 mm, totalizing approximately 30 follicles per routine. The follicles were dissected using tweezers, scissors and sterile scalpels. Dissected follicles were placed individually in 24 well-plates containing sterile saline solution and neatly ruptured with a pair of 18-G needles (1.2 x 40mm) to recover the cumulus-oocyte-complexes (COCs) and granulosa cells. Once located, COCs were classified and transferred individually to 20μl drops of handling (H) medium consisting of 25 mM HEPES-buffered Tissue Culture Medium-199 (TCM-199; GIBCO, Grand Island, NY, EUA) supplemented with 10% FBS (GIBCO), 0.2 mM sodium pyruvate, and 50 μg/mL gentamicin. The GCs samples belonging to each respective COCs were also individually collected with proper tracking code numbers, frozen in liquid nitrogen, and stored at -80 °C for further use. The classification of COCs was made according to the number of cumulus cell layers and cytoplasmic morphology in Grade I (GI), Grade II (GII), Grade III (GIII), or Grade IV (GIV), as previously described (Blondin & Sirard, 1995). However, only oocytes considered Grade I were used in the experiment.

### In vitro Maturation

COCs were cultured individually in 10 μL droplets of TCM-199 containing 10% FBS (Gibco Thermo Fisher, Massachusets - USA), 0.2 mM sodium pyruvate (Gibco Thermo Fisher, Massachusets - USA), 0.5 μg/mL follicle-stimulating hormone (Folltropin-V; Bioniche Animal Health Belleville, Canada), 5 U/mL, human Chorionic Gonadotropin-hCG (Chorulon®, Intervet International B. V., Boxmeer, Netherland) and 50 μg/mL gentamicin (Gibco Thermo Fisher, Massachusets - USA). Culture droplets were covered with mineral oil and placed in an incubator at 38.5°C and a humid atmosphere of 5% CO2 in atmospheric air. After 18 h of *in vitro* maturation, COCs were individually denuded of cumulus cells by 5 min exposure to a pre-warmed solution of trypsin (TrypLETE Express), followed by stripping with the aid of a 135mm diameter glass pipette. After denudation, oocytes with the presence of the first polar body (1PB), i.e., that completed the first meiotic division and arrested at MII, were returned to their respective maturation droplets for another two hours to complete a 20 h of in vitro maturation (IVM) period.

### Micromanipulation for Biopsy

After IVM, MII oocytes were exposed for 10 min to 10 ug/ml of HOESCHT 33342 in the maturation medium, washed in H medium, and placed in a biopsy dish containing 7.5ug/ml of cytochalasin under mineral oil. The microsurgical removal of the cytoplasmic biopsy was performed using an inverted microscope (Nikon Eclipse Ti) equipped with micromanipulators and microinjectors (Narishige, Tokyo, Japan). After alignment of first PB at 3h position, a biopsy of approximately 1% of the oocyte’s cytoplasm at 5h position was removed, verified for the absence of the MII spindle using UV light, and then transferred to a tube containing 1 U/uL RNase inhibitor (Sigma) in PBS. For the PB, it was placed in a tube containing only PBS and immediately frozen in liquid nitrogen. Then, Biopsied gametes were returned to their respective maturation droplet after five washes in the IVM medium until the end of oocyte maturation.

### Parthenogenetic activation and embryo culture

Biopsied oocytes were individually parthenogenetically activated to access their capacity to reach the blastocyst stage following standard procedures described in another study (Andrade et al., 2017). At 26 h of IVM, oocytes were washed in a 20 μL drop of H medium supplemented with 10% FBS, exposed to 5 μM of ionomycin, and washed in H medium with 30 mg/mL of BSA. After ionomycin treatment, oocytes were washed and ket for 3 h in synthetic oviduct fluid (SOF) supplemented with 5 mg/mL BSA, 2.5% FBS, 0.2 mM pyruvate, and 10 mg/ml of gentamicin and 2 mM of 6-dimethylaminopurine (6-DMAP). After 6-DMAP treatment, oocytes were washed and cultured individually for 8 days at 38.5 °C in 10 µl droplet of SOF medium in the presence of cumulus cells monolayer in an atmosphere of 5% O2, 5% CO2, and 90% N2. Activated oocytes were evaluated morphologically on D3 for cleavage and D7 for blastocyst rates. Morphological classification of cleavage rate on D3 was based on the presence of 8 symmetric blastomeres and the presence or not of a fully expanded or hatched blastocyst on D7. Finally, two groups of oocyte biopsy and granulosa samples were classified retrospectively according to whether the biopsied oocyte produced (BL) or not (CL) a blastocyst stage embryo at D7 of in vitro culture.

### RNA-Seq on granulosa cells and oocyte biopsies

A total of 5 routines were performed using follicles of 4 to 5 mm diameter, only follicles with oocytes classified as grade 1 were further utilized. Two of the best routines (higher partenote rates) were used for further sequencing. Granulosa cell samples RNA extraction was performed using the PicoPure RNA kit (ThermoFisher Scientific). RNA (1 ng) was used for cDNA synthesis and amplification using the SMART-Seq HT RNA-Seq library amplification kit (Takara Bio). According to the manufacturer’s recommendations, oocyte biopsy samples were directly converted to cDNA using the SMART-Seq kit (Takara). A total of eight (n = 8) biopsies were sequenced using a single-cell approach. A fraction of each sample was used for quality analysis using BioAnalyzer (Agilent Technologies, Wilmington, DE, USA). Thus, four (n = 4) biopsy and granulosa cell samples were sequenced from the embryos reaching blastocyst (BL) stage and arrested embryos at cleavage (CL) stage. Libraries were prepared using Nextera XT DNA Library Prep (Illumina) and sequenced on NextSeq2000 (Illumina, San Diego, USA) with 100 bp end reads. Reads were mapped to the bovine genome reference Bos taurus ARS-UCS1.2 using STAR and counting with Rsubread (version 1.34.7) (Dobin et al., 2012; Liao et al., 2019) with default settings, considering reads with sequencing scores across the total read length that were above 33. According to the ENSEMB, NCBI and NONCODE, RefSeq annotation, mapped reads were summarized at the gene level (Cunningham et al., 2022; Sayers et al., 2022; Zhao et al., 2021). This method is better described in our methodology study entitled as “**Assessment of Total Oocyte Transcripts Representation through Single Ooplasm Biopsy in bovine with High Reliability**”. Reads were filtered using the edgeR default threshold (Chen et al., 2020), where genes with more than three reading counts in at least three samples were used in DESeq2 model. Count summaries were obtained with the ’featureCounts’ function implemented in Rsubread Package with the strand-specific option and default settings. Differential gene expression analysis was performed using the DESeq2 package (Love et al., 2014). The Benjamin-Hochberg method “BH” with an alpha of 0.1 was used to compare the two groups, i.e., BL and CL. Enrichment analyses were performed using the ClusterProfiler R package software - Gene (Yu et al., 2012) Ontology [GO] - for biological process [bp], cell component [cc] and molecular function [mf], and the Encyclopedia of ’Kyoto’ Genes and Genomes [KEGG] considering BH (FDR < 0.05) (Kanehisa, 2000). PCA (Principal Component Analysis), smear plots (ggplot2), heatmaps (pheatmap), and pathways were performed and constructed using R (Kolde, 2019; Wickham, 2011). Package CeTF was used to create graphics for different RNA classes available in ENSEMBL annotation and transcription factors (Oliveira de Biagi Junior et al., 2021). Complementary analysis using SOTA – Self-organizing tree algorithm (clValid) was performed using the Euclidean distance and default parameters (Brock et al., 2008). Additional packages were used to customize graphics and for data manipulation: dplyr, tidyr (Wickham et al., 2019), fastqcr (Kassambara, 2018), tibble (Müller & Wickham, 2023), SummarizedExperiment (Morgan et al., 2022), enrichplot (Yu, 2023), rtracklayer (Lawrence et al., 2009), BiocGenerics (Huber et al., 2015), janitor (Firke et al., 2023), flextable (Gohel, 2023), grDevices (R Core Team, 2023), gplots (R. Warnes, 2022), scales (Wickham & Seidel, 2022), yulab.utils (Yu, 2022), ggdark (Grantham, 2019), pathview (Luo & Brouwer, 2013), png (Urbanek, 2022), ggrepel (Slowikowski, 2023), KEGGREST (Tenenbaum, 2022), agricolae (de Mendiburu, 2019), reshape2 (Wickham, 2020), readr (Wickham et al., 2023), graphics (R Core Team, 2023), googlesheets4 (Wickham et al., 2019), VennDiagram (Chen, 2022), GGally (Schloerke et al., 2021) and org.Bt.eg.db (Carlson, 2022).

### Single polar body WGBS

For bisulfite treatment of a single polar body, we followed the Wei protocol (Wei et al., 2019) and used the EpiTec Fast LyseAll Bisulfite Kit (Qiagen) with all volumes quartered. We added DNA protect buffer containing tetrahydrofurfuryl alcohol and then purified the sample, followed by water elution in 20 uL. The bisulfite-treated DNA fragments were ligated with a nine-nucleotide adapter and amplified using a GenomePlex Single Cell Whole Genome Amplification Kit (Sigma) for three rounds. We omitted the fragmentation step and amplified the samples for 15 rounds using the GenomePlex WGA Reamplification Kit (Sigma). We purified the samples using the QIAquick PCR Purification Kit and eluted them in 30 uL, then quantified them using a nanodrop. We ran the samples in an agarose gel to detect the fragment size, which resulted in fragments with ∼250 base pairs. We then sequenced the samples at the Nanuq facility - Centre d’expertise et de services Génome Québec, and after quality check, we prepared the library using the NEBNext® Ultra™ II DNA Library Prep Kit for Illumina®. We performed sequencing with a 10X coverage and concatenated the Fastq files in accordance with samples and strand. We verified the sequencing quality using the fastqcr package (Kassambara, 2018). We used MultiqQC (Ewels et al., 2016) to verify the results and applied the trimming process using Trim Galore, removing fragments < 30bp and with flags - q 24 --phred33 --length 30 --paired. We then performed fastqc followed by MultiQC to check the trimming process. We aligned the samples using BSMAP using Bos taurus ARS-UCD1.2 with flags -b -p 10 -w 100 -v 10. We performed peak calling using MACS3 with flags -t -n --fix-bimodal --nolambda --keep-dup all --extsize 100 -g hs -q 0.05 --cutoff-analysis (Zhang et al., 2008). We removed duplicates using samtools flagstat after sorting and imported *.bed files, separated peaks, and analyzed them using ChIPpeakAnno (Zhu et al., 2010). We found overlapped peaks using the findOverlapsOfPeaks function with maxgap = 1000 and connectedPeaks = “min”. Using the annotatePeakInBatch function with arguments output = “nearestBiDirectionalPromoters” and bindingRegion = c(-5000, 3000), we identified annotated peaks with gene names.

